# Using Community Science to Reveal the Global Chemogeography of River Metabolomes

**DOI:** 10.1101/2020.11.02.362905

**Authors:** Vanessa A. Garayburu-Caruso, Robert E. Danczak, James C. Stegen, Lupita Renteria, Marcy Mccall, Amy E. Goldman, Rosalie K. Chu, Jason Toyoda, Charles T. Resch, Joshua M. Torgeson, Jacqueline Wells, Sarah Fansler, Swatantar Kumar, Emily B. Graham

## Abstract

River corridor metabolomes reflect organic matter (OM) processing that drives aquatic biogeochemical cycles. Recent work highlights the power of ultrahigh-resolution mass spectrometry for understanding metabolome composition and river corridor metabolism. However, there have been no studies on the global chemogeography of surface water and sediment metabolomes using ultrahigh-resolution techniques. Here, we describe a community science effort from the Worldwide Hydrobiogeochemistry Observation Network for Dynamic River Systems (WHONDRS) consortium to characterize global metabolomes in surface water and sediment that span multiple stream orders and biomes. We describe the distribution of key aspects of metabolomes including elemental groups, chemical classes, indices, and inferred biochemical transformations. We show that metabolomes significantly differ across surface water and sediment and that surface water metabolomes are more rich and variable. We also use inferred biochemical transformations to identify core metabolic processes shared among surface water and sediment. Finally, we observe significant spatial variation in sediment metabolites between rivers in the eastern and western portions of the contiguous United States. Our work not only provides a basis for understanding global patterns in river corridor biogeochemical cycles but also demonstrates that community science endeavors can enable global research projects that are unfeasible with traditional research models.

## 1. Introduction

Organic matter (OM) transformations in aquatic ecosystems are a critical source of uncertainty in global biogeochemical cycles [1–4]. More than half of OM inputs to freshwater ecosystems are metabolized before reaching the oceans [1,2,4], yet while several studies have focused on quantifying OM uptake and export rates [1,5,6], the processes driving river corridor OM transformations across spatial scales remain poorly understood.

River corridor OM pools contain an extensive variety of molecules that are both produced and metabolized by microorganisms, which are processes reflected in the composition of sediment and surface water metabolomes [2,7,8]. Metabolic transformations of OM in freshwater ecosystems have been traditionally estimated by a combination of laboratory incubations and in-stream tracer additions [9–12]. However, results from incubation experiments are challenging to scale beyond laboratory conditions [9,10], and in-stream tracer processing often does not reflect ambient biogeochemical processes, as the naturally occurring metabolome is more chemically diverse than the tracer added to the stream [11,12]. Several studies have shown that OM pool composition can influence microbial activity, highlighting complexities in the metabolic processes that determine OM transformations [13–18]. Consequently, determining mechanisms underlying river corridor metabolome composition at a large scale remains challenging.

Environmental metabolomics uses the identification of small molecules in an organism (metabolites) to characterize the interactions of organisms within their environment [19]. Over the past several years, this definition has been extended to encompass all metabolites present in complex environmental systems for which it is difficult to attribute specific metabolites to specific organisms [20–24]. Different metabolomic techniques have been implemented across fields to enhance our understanding of microbial communities [25,26], anthropogenic activities and pollution sources [27–29], and potential bioremediation strategies [30]. Recently, environmental metabolomics, enabled by ultrahigh-resolution mass spectrometry, has allowed us to reveal connections between OM character, reactivity, and biochemical transformations within and across river ecosystems [15,17,18,31–35]. These advances have vastly improved our understanding of the mechanisms governing OM bioavailability and biochemical transformations at a global scale. For instance, previous studies have used ultrahigh-resolution metabolomics from river water across different climatic regions to find common compositional features that would inform global carbon dynamics [36] and to investigate environmental drivers affecting OM composition, bioavailability, and transport of OM [37]. In addition, recent studies show that OM thermodynamics influence aerobic respiration under carbon-limited scenarios [16], that biogeochemical hotspots are influenced by OM nitrogen content [17], and that hyporheic zone mixing induces OM metabolism via a priming effect [15]. These detailed metabolome characterizations have the potential to enable global-scale inferences about watershed features (e.g., vegetation, lithology, hydrology, microbiology, climate) that govern the reactivity and fate of OM across river corridors [35,38]. In turn, metabolomics can enhance our predictive capabilities of global river corridor biogeochemical cycles by helping to improve the representation of biochemical mechanisms in numerical models, such as reactive transport codes [39,40]. For example, an emerging substrate-explicit model uses thermodynamic theory to explicitly account for the chemical composition of all metabolites in OM pools to improve the predictive capacity of biogeochemical models [40].

Characterizing metabolomes across global spatiotemporal scales requires a way to collect multiple data types across diverse locations in such a way that they can be analyzed together. This goal can be facilitated by a framework that requires studies to Integrate biological, physical, and chemical processes across scales; Coordinate with consistent methods; be Open across the research lifecycle; and Network with global collaborators to reduce the burden on a single team (ICON) [41,42]. When ICON principles are applied, they allow for distributed sampling in ways that have historically been difficult to achieve.

The Worldwide Hydrobiogeochemistry Observation Network for Dynamic River Systems (WHONDRS) is a global consortium of researchers based out of Pacific Northwest National Laboratory that uses an ICON-based approach to understand coupled hydrologic, biogeochemical, and microbial functions in river corridors [35]. ICON principles allow WHONDRS to collect open, globally distributed data through collaboration with the scientific community. The WHONDRS consortium designs sampling campaigns that target specific spatial and temporal scales, modifies its approach based on community input, and then sends free sampling kits to collaborators. All WHONDRS data are openly accessible through Environmental Systems Science Data Infrastructure for a Virtual Ecosystem (ESS-DIVE-https://data.ess-dive.lbl.gov/) and the National Center for Biotechnology Information (NCBI), and the WHONDRS consortium ascribes to FAIR data principles (findable, accessible, interoperable, reusable) [43]. This approach enables WHONDRS to collect, analyze, and distribute ultrahigh-resolution metabolomic data to the global scientific community.

Here, we describe a community science effort conducted by the WHONDRS consortium during July-August 2019 that used Fourier-transform ion cyclotron resonance mass spectrometry (FTICR-MS) to characterize metabolomes in global surface water and sediment spanning a range of biomes (e.g., desert-like in the Columbia Plateau, subtropical in southern Florida, temperate forests in the Mid-Atlantic) and stream orders [44]. We describe key metabolome characteristics of surface water and sediment and also explore spatial variation of these characteristics within the United States. We focus on central aspects of metabolomes including assigned elemental groups, chemical classes, descriptor indices, and biochemical transformations. This paper provides a benchmark for studying integrated surface water and sediment river corridor metabolomes and highlights the need to engage a wider scientific community in order to expand the reach and impact of scientific advancements.

## 2. Results and Discussion

### 2.1. Surface Water Metabolome is More Unsaturated, Aromatic, Oxidized, Rich, and Variable than Sediment Metabolome

In order to assess patterns in global metabolome composition, we derived a number of descriptive metrics that summarize FTICR-MS metabolomic profiles. Specifically, we compared double-bond equivalents (DBE), modified aromaticity index (AI_Mod_), nominal oxidation state of carbon (NOSC), inferred chemical classes (e.g., lignin-like, protein-like), and elemental groups (e.g., CHO, CHON, CHOSP) of surface water and sediment metabolomes. The double-bond equivalent metric (DBE) describes the degree of chemical unsaturation of bonds in a particular metabolite [45,46], AI_Mod_ quantifies the degree of aromaticity (i.e., ring-like shape) of a metabolite [45–47], and NOSC indicates the energy required to oxidize different metabolomes [48]. High values of AI_Mod_ can denote the existence of either aromatic (AI_Mod_ > 0.5) or condensed aromatic structures (AI_Mod_ ≥ 0.67), and high DBE indicates more saturated compounds. NOSC is inversely correlated with the Gibbs free energy of carbon oxidation. Higher NOSC corresponds to metabolites that are more oxidized and thermodynamically favorable [15–18,48,49]. Chemical class assignments for each metabolite were predicted using oxygen-to-carbon and hydrogen-to-carbon ratios (i.e., Van Krevelen classes [50]). Finally, we used the molecular formula assigned to each metabolite to describe the relative abundance of different heteroatom combinations associated with CHO groups (i.e., differences in −N, −S and/or −P). We then compared metrics across all metabolites found in any surface water sample vs. all metabolites found in any sediment sample. All analyses in Section 2.1 were conducted only on FTICR-MS peaks that were able to be assigned a molecular formula. Other metrics describing metabolome composition are reported in the SI (Table S1).

Surface water metabolomes were composed of comparatively more unsaturated and aromatic compounds with a higher nominal oxidation state than sediment. This was denoted by significantly higher AI_Mod_, DBE, and NOSC than sediment metabolomes (Figure 1, *p*-value < 0.001). In addition, we observed higher relative abundances of lignin-like, tannin-like, and condensed-hydrocarbon-like metabolites in surface water versus sediment (Figure 2, all *p*-values < 0.001). These classes of metabolites are characteristic of terrestrial OM [51], and their prevalence in surface water metabolomes indicates a larger contribution of terrestrial OM in surface water relative to sediment. This may also indicate greater contributions of microbially processed OM in sediment, as has been observed previously in comparisons between surface water and hyporheic zone porewater [15, 52].

**Figure 1.**
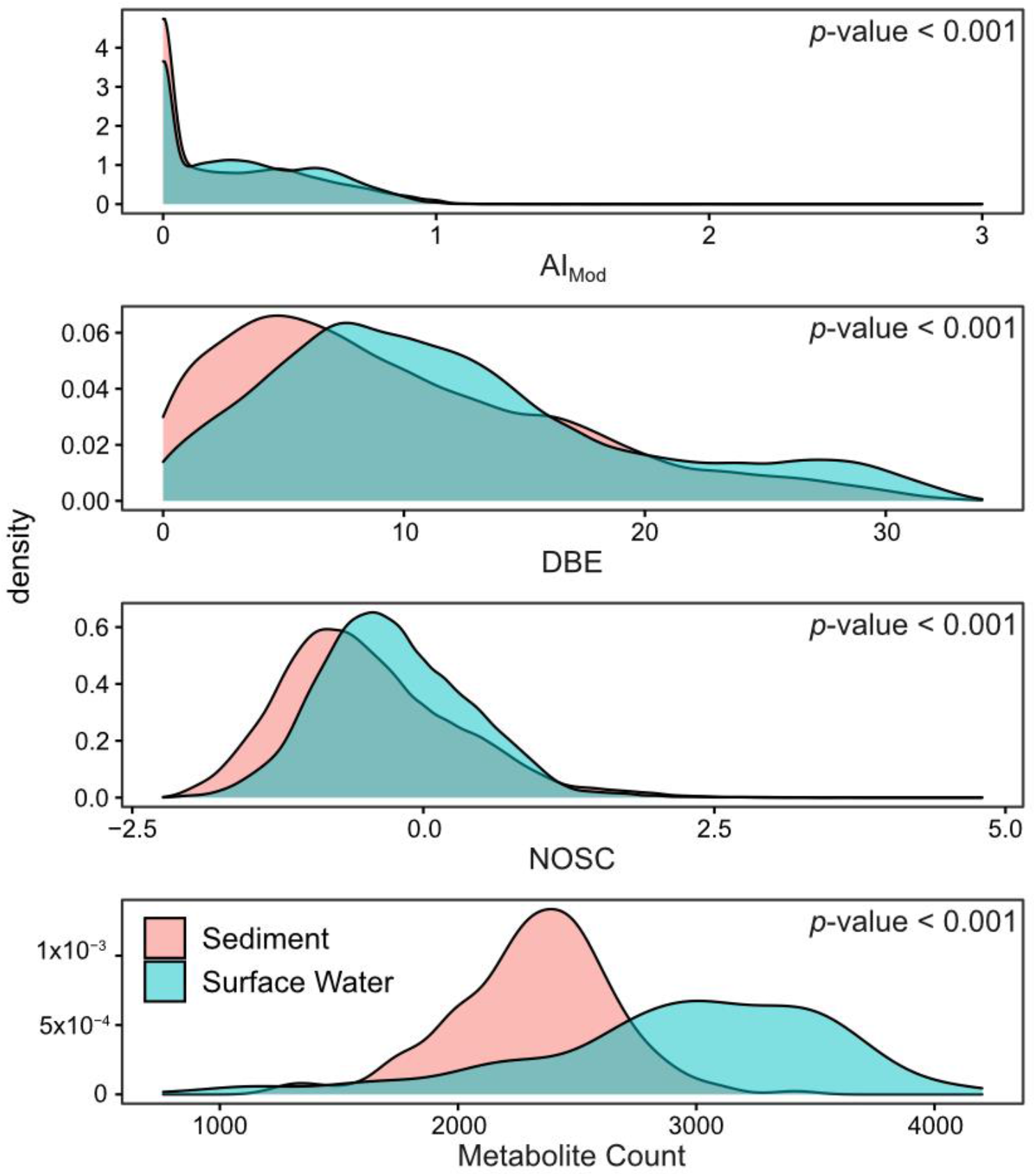
Density plots comparing the properties of all molecular formulas found in surface water and sediment samples. AI_Mod_ (modified aromaticity index) is a measure of the potential ring-like structure in a given molecular formula. DBE (double-bond equivalents) is an approximation of potential unsaturation. NOSC (nominal oxidation state of carbon) represents the degree of oxidation/reduction of a given molecular formula. Significance values obtained via a two-sided Mann–Whitney U test to compare sample type distributions are denoted in the upper right corner of each panel.

**Figure 2.**
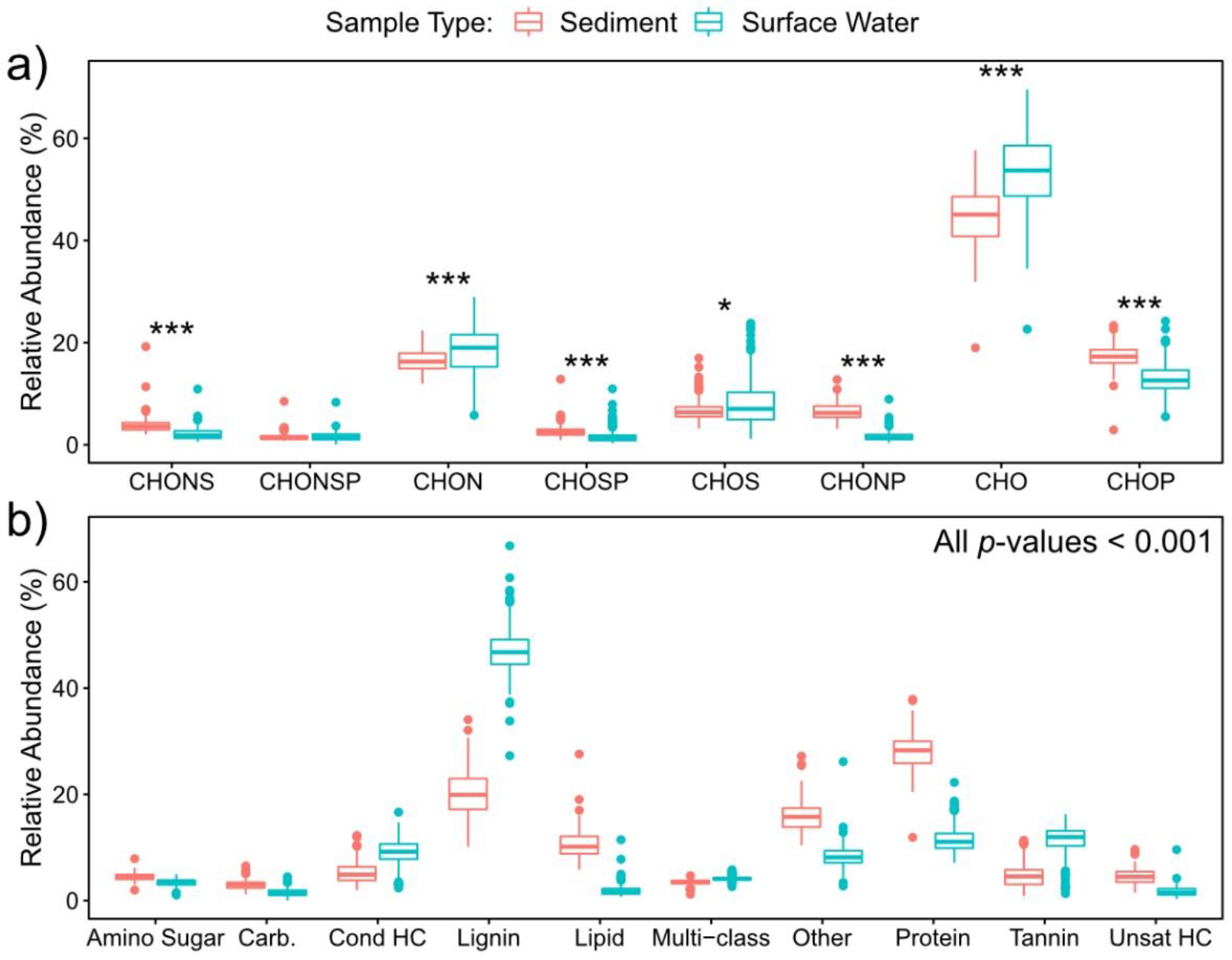
Box plots comparing the relative abundances of metabolites belonging to specific elemental groups (**a**) and chemical classes (**b**) between sediment and surface water. As in Figure 1, these values were obtained from all metabolites assigned molecular formulas in sediment and surface water. Significance values were obtained via a two-sided Mann–Whitney U test (0.05 > * > 0.01 > ** > 0.001 > ***).

The higher relative abundance of unsaturated and aromatic metabolites in surface water contrasts with previous studies that have observed that these compounds are more common in sediment porewater than in lake surface water or aquifer recharge water [53,54]. These studies inferred low physical, chemical, and/or biological transformation of sediment porewater associated OM. This deviation might be connected to the systems studied. For example, Pracht et al. [53] examined a system where sediment OM was protected due to physical and/or chemical constraints such as mineral sorption and hydrophobic encapsulation [53]. We studied rivers where the shallow benthic layer and the hyporheic zone are known to enhance biogeochemical reactions [55–59]. In turn, we hypothesize that very high rates of biological activity in riverbed sediment [60,61] could be responsible for lower AI_Mod_, DBE, and NOSC values of sediment metabolomes relative to surface water, in contrast to previous work in potentially less active lake and aquifer systems.

In addition, the relative abundances of lipid-like and protein-like metabolites were significantly higher in sediment than in surface water (*p*-value < 0.001). More lipid-like compounds in sediment could reflect higher microbial biomass [62] and further supports our inference that sediment metabolomes were influenced by microbial processes to a greater extent than surface water metabolomes. This highlights the key role played by riverbed sediment and associated hyporheic zones in river corridor biogeochemistry that likely influences global elemental cycles but is not captured in current Earth system models.

Conversely, elemental groups of metabolites were similar across surface water and sediment. The median abundance of each elemental group did not vary more than 5% between the two environments, except for CHO (~9%) and CHONSP (~0.2%, not statistically significant) groups. Bulk similarities in elemental groups, in contrast to chemical classes, could indicate that the presence or absence of heteroatoms alone is insufficient to distinguish metabolomes and that elemental stoichiometry of the entire metabolite (the basis for chemical class assignment) may be more important for distinguishing metabolomes. This is important because the elemental stoichiometry of metabolites can mechanistically connect OM thermodynamics to biogeochemical reactions and rates [40].

Metabolites found in surface water were distinct from and showed more among-sample variation than those in sediment. To evaluate compositional differences, we conducted a principal component analysis (PCA) and a beta-dispersion analysis (Figure 3). For consistency with prior analyses in this section, we present a PCA on only peaks assigned a molecular formula in Figure 3. When performed on all peaks, regardless of formula assignment, PCA results were consistent with Figure 3 (Figure S1, Table S2). The PCA, in conjunction with a PERMANOVA comparison, indicated that surface water and sediment metabolomes significantly diverged in composition (*p*-value < 0.001). Loadings for PC1 and PC2 are presented in Table S3. In general, the loadings suggest that many metabolites contributed to the separation between surface water and sediment metabolomes (PC1), while CHON-containing metabolites primarily drove variability in surface water metabolomes (PC2). The beta-dispersion analysis further indicated that surface water metabolomes were more dispersed in multivariate space than sediment metabolomes (Figure 3; *p*-value < 0.001). Additionally, surface water metabolomes had higher richness (i.e., more peaks with assigned formulas detected on average) than sediment metabolomes (Figure 1). These patterns indicate that metabolomes in surface water and sediment may be shaped by distinct processes that likely span differences in inputs, rates of microbial activity, and abiotic constraints.

**Figure 3.**
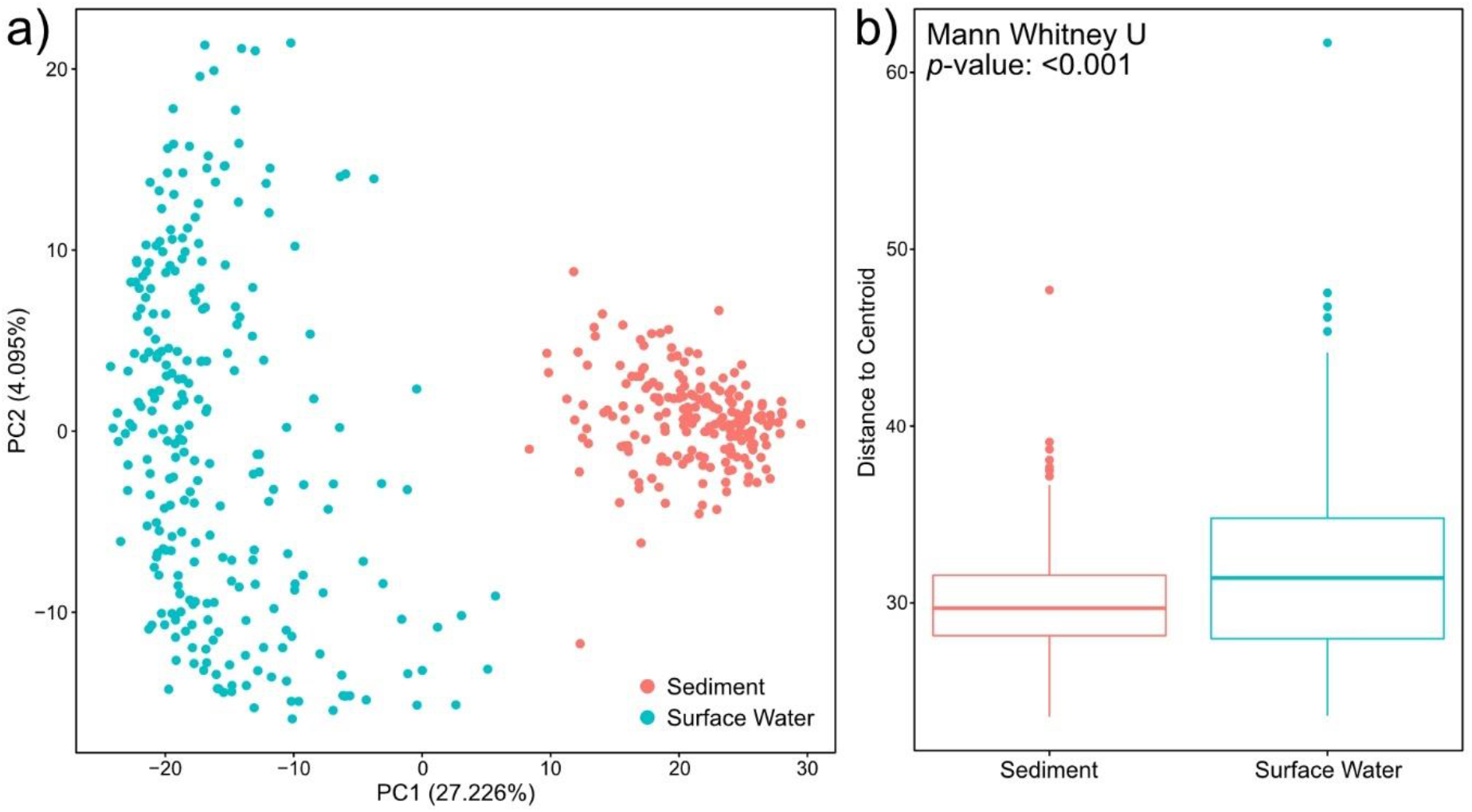
Principal component analysis (PCA) of the molecular formula data (**a**). Differences between surface water and sediment metabolomes were significant per a Euclidean distance-based PERMANOVA (*p*-value < 0.001). The degree of among-sample variation was evaluated by quantifying beta-dispersion. Surface water had higher beta-dispersion per a two-sided Mann– Whitney U test (*p*-value < 0.001) (**b**).

We hypothesize that higher richness and greater among-sample variation in surface water metabolomes could reflect more heterogeneous environmental pressures. For example, we sampled across a broad range of latitudes and stream orders that likely led to among-site variation in light exposure (Table S4 [44]). This may, in turn, have led to variation in surface water temperatures and surface water metabolite photodegradation, thereby increasing metabolome variability and richness [63]. Hydrology could also contribute to metabolome richness in surface water as precipitation events and associated runoff transport large amounts of terrestrial OM into rivers [64]. For example, precipitation has been shown to increase aromatic OM and decrease more labile OM in surface water [65–67]. We hypothesize that more immediate connectivity between surface water and terrestrial systems, relative to connectivity between sediment and terrestrial systems, is at least partially responsible for greater variation and higher richness in surface water metabolomes.

In addition, lower sediment metabolome variation and richness could be due to comparatively higher rates of microbial activity in sediment that degrade polymeric OM into a limited set of less chemically complex metabolites. This would result in a reduction in the number of distinct metabolites present by collapsing a diverse pool of OM into microbial exudates. Sediment metabolomes may also be constrained by interactions with sediment mineral surfaces, especially considering that we studied only the water-extractable metabolome. This subset of the full sediment metabolome may inherently be composed of a restricted suite of metabolites [68], leading to lower among-sample variation and lower richness. We nonetheless hypothesize that among-site variation in mineralogy could contribute to some of the observed sediment metabolome variation. The WHONDRS consortium is currently generating mineralogy data to test this hypothesis.

Together, the observations presented in this section indicate that there are significant differences across global surface water and sediment metabolomes, where surface water metabolomes are more unsaturated, aromatic, oxidized, rich, and variable. These characteristics suggest that surface water metabolomes are more dynamic due to a variety of watershed and river corridor processes (discussed above), while sediment metabolomes may be more stable integrators of localized processes (e.g., mineral interactions and microbial processing of OM).

### 2.2. Nitrogen-, Sulfur- and Phosphorous-Containing Transformations Vary Across Surface Water and Sediment Metabolomes

We evaluated how potential reactions in metabolomes varied across the globe by inferring biochemical transformations as per Bailey et al. [62], Kaling et al. [69], Moritz et al. [70], Graham et al. [17,18], Garayburu-Caruso et al. [16], Danczak et al. [38], and Stegen et al. [15]. This method leverages the ultrahigh-resolution of FTICR-MS to compare mass differences between detected peaks to a database of common biochemical transformations. Identified biochemical transformations provide information regarding the frequency at which a specific molecule could have been gained or lost during metabolism. Resulting transformation counts can then be separated based upon their chemical properties to study the potential role of the molecule gained or lost in metabolome composition. Unlike the analyses described in the previous section, where formula assignments of metabolites are necessary, this method allows for the incorporation of all detected metabolites into downstream analyses.

We observed that biochemical transformations involving molecules containing nitrogen (N), sulfur (S), or phosphorous (P) exhibited divergent patterns between surface water and sediment metabolomes (Figure 4). Specifically, surface water had a significantly higher relative abundance of N-containing transformations, while sediment metabolomes had more S- and P-containing transformations (*p*-value < 0.001 in all cases). This contrast between surface water and sediment may occur due to variation in nutrient requirements within the water column and sediment, as biochemical transformations have been inferred to reflect nutrient limitations in other systems. For example, Garayburu-Caruso et al. [16] found an increase in N-containing transformations under nutrient-limited conditions, but only when N-containing OM was introduced. More N-containing transformations in surface water as compared to sediment could therefore reflect microbial N mining in surface water through the preferential decomposition of N-containing OM [71,72]. In addition, a higher abundance of P- and S-containing biochemical transformations in sediment further suggest that microbial metabolism is limited by different factors between surface water and sediment environments, which are potentially associated with nutrient assimilation processes [73,74].

**Figure 4.**
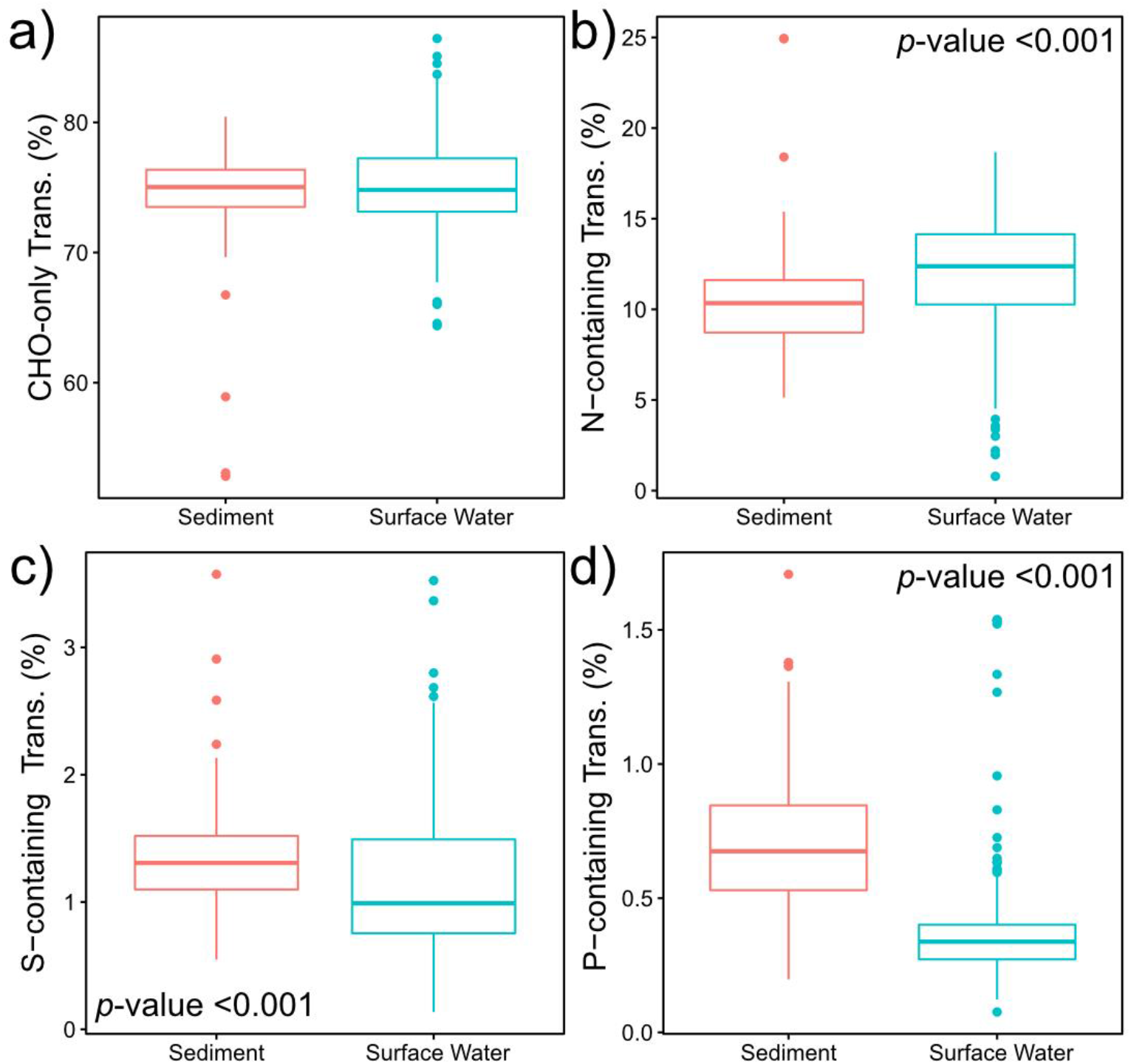
Boxplots displaying the patterns of CHO-only transformations (**a**); N-containing transformations (**b**); S-containing transformations (**c**); and P-containing transformations (**d**). False discovery rate (FDR)-corrected two-sided Mann-Whitney U test *p*-values are provided in either the top right or the bottom left corner of panels with significant comparisons.

Interestingly, the higher abundance of N-containing transformations in surface water observed in this study is contrary to past work, where porewater had a higher relative abundance of N-containing transformations than surface water [15,38]. Given that these studies were collected from the Pacific Northwest region of the United States (e.g., eastern Washington and Oregon), this highlights the necessity to expand research beyond individual test systems or geographic regions and emphasizes the utility of global studies through efforts like WHONDRS.

In contrast, biochemical transformations that did not involve N-, S-, or P-containing molecules were not significantly different between surface water and sediment metabolomes. Similar patterns have been observed in other studies [38]. These results indicate the presence of ubiquitous biochemical transformations that occur in both surface water and sediment (Figure 4). Based on these results, we hypothesize that N-, S-, and P-containing transformations may have a stronger dependency than CHO-only transformations on nutrient status. That is, changes in nutrient availability across surface water and sediment environments may drive shifts in N-, S-, and/or P-containing transformations but not influence transformations that do not involve these nutrients. Additional data on variation in nutrient limitation and availability will be required to test this hypothesis.

### 2.3. Sediment Metabolomes are More Spatially Variable Than Surface Water Metabolomes

Because most of our sampling locations were in the contiguous United States (CONUS), we used CONUS data to resolve potential spatial patterns in metabolomes (Figures 5 and 6). In order to uncover site-by-site metabolomic variation, we calculated the mean value for each derived metric (e.g., AI_Mod_, NOSC, DBE) and calculated the relative abundance of elemental groups and chemical classes (e.g., CHO and lignin-like) for each sample.

**Figure 5.**
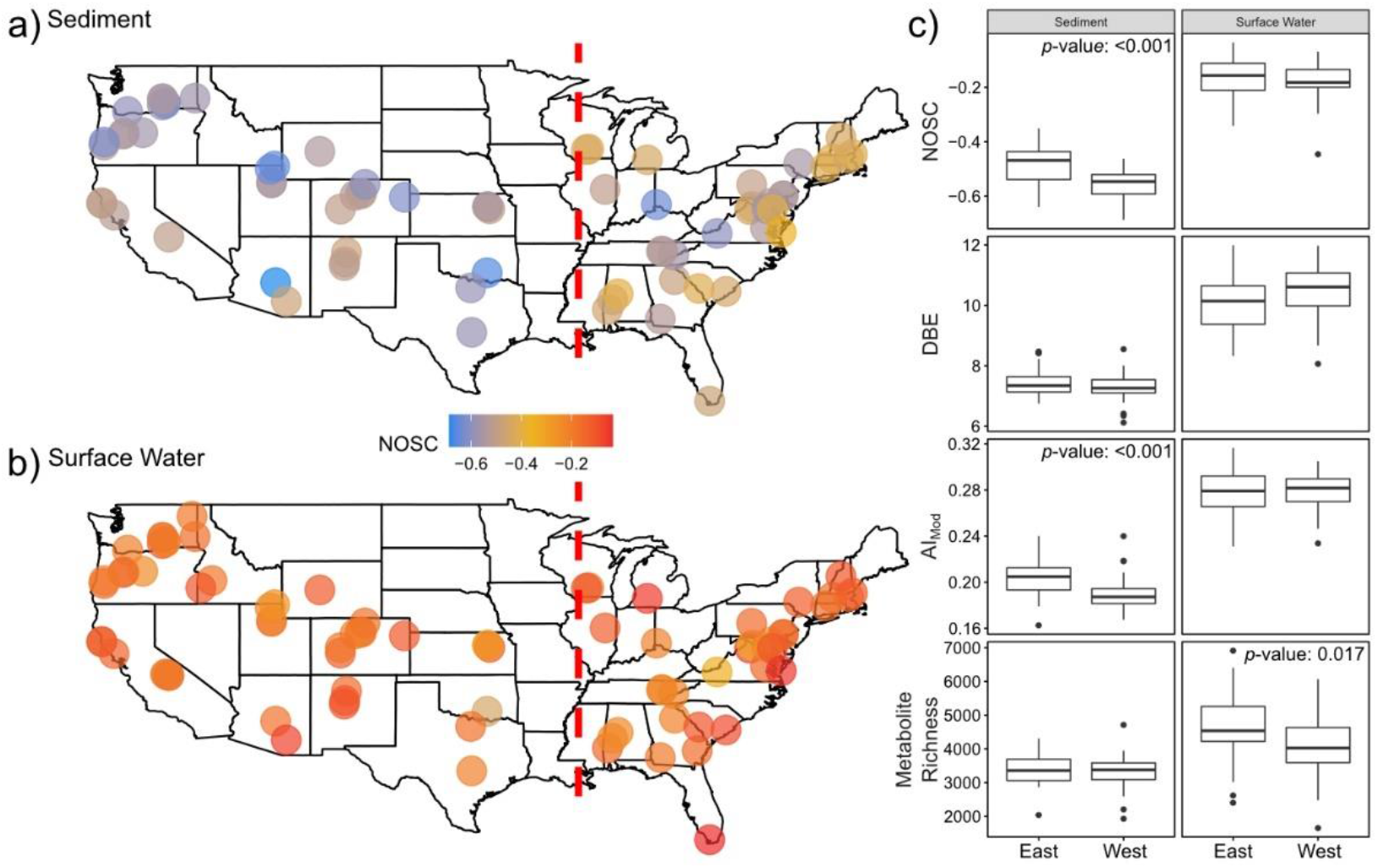
Maps of the United States revealing the spatial variability of average NOSC in sediment (**a**) and surface water (**b**) metabolomes. East vs. West spatial patterns for various derived metrics are displayed in panel (**c**). Statistically significant differences identified via a two-sided Mann–Whitney U test are indicated by *p*-values listed in each comparison. The red dashed line represents the longitude of the Mississippi River at St. Louis, MO, USA—the dividing line between East and West samples.

**Figure 6.**
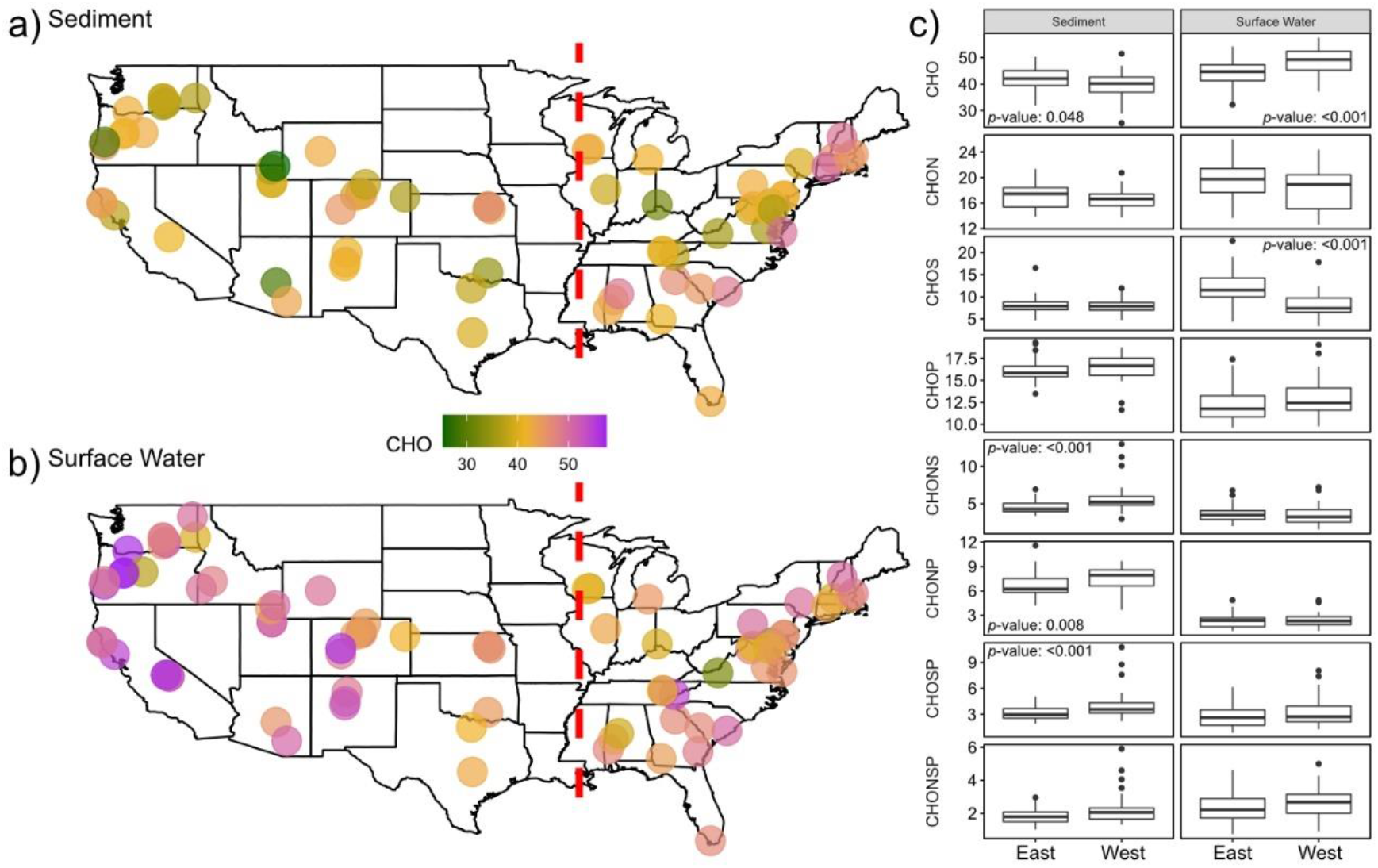
Maps of the United States revealing the spatial variability of the relative abundance of metabolites containing only CHO with sediment (**a**) and surface water (**b**); East vs. West spatial patterns for different elemental groups’ relative abundances are displayed in panel (**c**). Statistically significant differences identified via a two-sided Mann–Whitney U test are indicated by *p*-values listed in each comparison. The red dashed line represents the longitude of the Mississippi River at St. Louis, MO, USA—the dividing line between East and West samples.

Overall, differences between the mean properties of CONUS surface water and sediment metabolomes were generally consistent with differences between global surface water and sediment metabolomes reported in Figure 1. For instance, surface water metabolomes displayed higher AI_Mod_, DBE, and NOSC than sediment metabolomes (Figure 5, *p*-value < 0.001 for all, Table S5). We also observed similar patterns in both the relative abundances of specific elemental groups (Figure 6, *p*-value < 0.001 for all, Table S5) and chemical classes (*p*-value < 0.001 for all, Table S5 and File S1). In order to expand our analyses, we investigated spatial patterns in individual metabolomic features across the CONUS by comparing sites that were east (hereafter “East”, surface water *n* = 34, sediment *n* = 33) vs. west (hereafter “West”, surface water n = 45, sediment n = 38) of the Mississippi River.

In general, we observed greater spatial patterning in sediment metabolomes than in surface water metabolomes. Average NOSC and AI_Mod_ values of sediment metabolomes were higher in the East than in the West (*p*-value < 0.001 for both). Metabolites containing CHONP, CHONS, or CHOSP constituted a significantly lower relative abundance of metabolomes in the East sediment, relative to the West (*p*-value < 0.001 for all). Lastly, lignin- and tannin-like metabolites constituted a higher relative abundance of sediment metabolomes in the East, while protein-, condensed-hydrocarbon-, and unsaturated-hydrocarbon-like metabolites were more abundant in the West (*p*-value < 0.001 for all, Table S5 and File S1).

In contrast to spatial patterns in sediment metabolomes, surface water showed less spatial structure with only CHO and CHOS metabolites showing significant shifts between East and West (*p*-value < 0.001 for both). CHO and CHOS metabolites comprised higher and lower relative abundances in the West than in the East, respectively (Figure 6). No spatial patterns were observed in NOSC (*p*-value = 0.4, Table S5), DBE (*p*-value = 0.07, Table S5), or AI_Mod_ (*p*-value = 0.93, Table S5) in surface water (Figure 5). Results from the remainder of molecular indices, elemental groups, and chemical class comparisons are shown in Table S5 and File S1.

Together, these patterns suggest that spatial differences in the metabolic processes driving OM cycling in the East versus the West have a stronger influence on sediment metabolomes than surface water metabolomes. While our non-spatial analyses showed greater among-sample and among-site variability in surface water metabolomes (Figure 3), the lack of spatial structure across the CONUS suggests that this variability is not driven by factors that are spatially structured at the continental scale. Instead, we hypothesize that surface water metabolomes are more temporally variable due to fluctuating inputs from precipitation events. To more directly evaluate spatial structure in surface water metabolomes, it is likely necessary to control for precipitation history and hydrologic connectivity to terrestrial systems. These inferences are supported by previous studies addressing spatial dissolved OM chemography dynamics showing that longitudinal patterns of dissolved OM in surface water are sensitive to hydrologic events [75–77]. However, there is a complex interaction between hydrology and space, as the sources and quality of OM from different regions may respond differently to hydrological variation [78].

The contrasting patterns in surface water and sediment metabolome characteristics across the East-West gradient could be the result of many factors, including vegetation cover [79], underlying lithology [80], photoreactivity [63], climate and precipitation regime [80], and/or microbial metabolism [81]. Additional data and analyses will be required to disentangle the relative contributions of these potential drivers. Pursuing this knowledge is important for explaining and ultimately predicting OM transformations. In turn, representing these processes and their impacts on biogeochemical cycles in processed-based models has the potential to improve the accuracy of biogeochemical predictions across the globe.

## 3. Materials and Methods

### 3.1. WHONDRS Summer 2019 Sampling Campaign

In July and August 2019, the WHONDRS consortium initiated a study of global river corridors to evaluate interactions between ecosystem features, microbial communities, and metabolomes in surface water and shallow sediments. To design the study, the WHONDRS consortium held multiple webinars with collaborators who volunteered to collect samples. The webinars allowed for community input on sampling protocol and data collected. More details are available at https://whondrs.pnnl.gov.

Briefly, WHONDRS developed sampling protocols and videos in coordination with the scientific community that were made openly available via YouTube, sent free sampling kits to collaborators, and conducted a suite of biogeochemical analyses on surface water and sediment. All data will be made open access following QA/QC at https://data.ess-dive.lbl.gov/. Preliminary data are available on a Google Drive linked via https://whondrs.pnnl.gov as they become available.

The 2019 study collected samples and metadata associated with stream order, climate, vegetation, and geomorphological features from 97 river corridors in 8 countries within a 6-week period, from 29 July to 19 September (Table S4, [44]). Stream order information (Table S4) was acquired for sites within the continental United States through the EPA National National Hydrography Dataset Plus (https://www.epa.gov/waterdata/nhdplus-national-hydrography-dataset-plus), and stream orders for a couple of Canada sites were acquired through British Columbia Data Catalogue (https://catalogue.data.gov.bc.ca/dataset/75299593-3222-40f9-879f-29e9824fc978). Stream orders indicate the relative size of a stream [82]. The data provided in the SI were calculated following Strahler’s definition of stream order [83]. This is estimated based on the size of its tributaries; for example, if two 1st order streams come together, they will form a 2nd order stream. Lower stream orders tend to be small tributaries or headwaters, while large stream orders are often major rivers [82,83].

This paper focuses on surface water (95 sites) and sediment (78 sites) collected across biomes (i.e., desert, tropical, temperate forests), from which a total of 504 samples were analyzed. Toyoda et al. [44] provide additional metadata associated with specific site characteristics (e.g., hydrogeomorphology, vegetation, temperature, discharge).

### 3.2. Sample Collection and Laboratory Pre-Processing

At each location, collaborators selected sampling sites within 100 m of a station that measured river discharge, height, or pressure. Within each site, 3 depositional zones were identified for sediment collection following NEON’s protocol (NEON.DOC.001193; [84]) and labeled as upstream, midstream, or downstream. The depositional zones were situated within 10 m of each other. Surface water was sampled in triplicate prior to sediment sampling. Surface water was collected only at the downstream site before collecting the sediments to make sure the water collected was not affected by sediment debris mobilized during water or sediment sampling at upstream locations. Sediments were collected from all three zones, where each zone provided a biological replicate of a sediment sample.

Surface water was collected using a 60 mL syringe and was filtered through a 0.22 μm sterivex filter (EMD Millipore) into a 40 mL glass vial (I-Chem amber VOA glass vials; ThermoFisher, pre-acidified with 10 μL of 85% phosphoric acid). Subsequently, 125 mL of surface sediments (1–3 cm depth) were sampled from a ~1m2 area at each depositional zone with a stainless steel scoop, making sure the sediments were saturated upon collection. All samples were shipped to Pacific Northwest National Laboratory on blue ice within 24 h of collection.

Surface water samples were immediately frozen at −20 °C upon receiving. Sediments from each depositional zone were individually sieved to <2 mm, subsampled into proteomic friendly tubes (Genesee Scientific), and stored at −20 °C for FTICR-MS analysis.

### 3.3. Fourier Transform Ion Cyclotron Resonance Mass Spectrometry (FTICR-MS)

Surface water samples were thawed in the dark at 4 °C for 72 h. Non-purgeable organic carbon (NPOC) was determined using a 5 mL aliquot of the acidified water sample by a Shimadzu combustion carbon analyzer TOC-L CSH/CSN E100V with ASI-L autosampler. NPOC concentrations (Table S6) were normalized to 1.5 mg C L⁻¹ across all samples to allow for data comparison across sites within this study and other WHONDRS sampling campaigns. Diluted samples were acidified to pH 2 with 85% phosphoric acid and extracted with PPL cartridges (Bond Elut), following Dittmar et al. [85].

Sediment samples were thawed overnight in the dark at 4 °C. Then, sediment organic matter was extracted in proteomic friendly tubes (Genesee Scientific) with a 1:2 ratio of sediment to water (5 g of sediment to 10 mL of milli-Q water). During the extraction, tubes were continuously shaken in the dark at 375 rpm and 21 °C for 2 h, after which the tubes were centrifuged at 6000 rcf and 21 °C for 5 min. The supernatant was collected and filtered through 0.22 μm polyethersulfone membrane filter (Millipore Sterivex, USA) into borosilicate glass vials. NPOC (Shimadzu combustion carbon analyzer TOC–Vcsh with ASI–V autosampler) was determined using a 5 mL aliquot from the filtered supernatant. As with the water samples, this supernatant was normalized to a standard NPOC concentration (Table S4) of 1.5 mg C L^−1^, acidified to pH 2 with 85% phosphoric acid, and extracted with PPL cartridges following the same methods described above.

A 12 Tesla (12 T) Bruker SolariX Fourier transform ion cyclotron mass spectrometer (FTICR-MS; Bruker, SolariX, Billerica, MA, USA) located at the Environmental Molecular Sciences Laboratory in Richland, WA, was used to collect ultrahigh-resolution mass spectra of surface water and sediment OM pools. Resolution was 220 K at 481.185 *m/z.* The FTICR-MS was outfitted with a standard electrospray ionization (ESI) source, and data were acquired in negative mode with the voltage set to +4.2 kV. The instrument was externally calibrated weekly to a mass accuracy of <0.1 ppm; in addition, the instrument settings were optimized by tuning on a Suwannee River Fulvic Acid (SRFA) standard. Data were collected with an ion accumulation of 0.05 sec for surface water and 0.1 or 0.2 sec for sediment from 100–900 *m/z* at 4 M. One hundred forty-four scans were co-added for each sample and internally calibrated using an OM homologous series separated by 14 Da (–CH2 groups). The mass measurement accuracy was typically within 1 ppm for singly charged ions across a broad *m/z* range (100 *m/z*–900 *m/z*). BrukerDaltonik Data Analysis (version 4.2) was used to convert raw spectra to a list of *m/z* values by applying the FTMS peak picker module with a signal-to-noise ratio (S/N) threshold set to 7 and absolute intensity threshold to the default value of 100. We aligned peaks (0.5 ppm threshold) and assigned chemical formulas using Formularity [86]. The Compound Identification Algorithm in Formularity was used with the following criteria: S/N > 7 and mass measurement error <0.5 ppm. This algorithm takes into consideration the presence of C, H, O, N, S, and P and excludes other elements.

It is important to note that FTICR-MS is not quantitative and does not provide information about the structure of the molecular formulas identified. This method provides a non-targeted approach to reliably identify molecular formulas of organic metabolites with masses between 200–900 *m/z*. The power of FTICR-MS is that it can capture thousands of metabolites simultaneously in contrast to other global environmental metabolomics techniques that yield less information. A key consideration with FTICR-MS-derived information is that it captures all ionizable organic molecules and thus is source-agnostic (e.g., not all detected compounds are guaranteed to be biologically derived). Hence there is a tradeoff of depth vs. specificity in metabolomics methods, and FTICR-MS sacrifices some specificity for depth. In addition, the sediment-water extractions performed in this study provide chemical selectivity towards water-extractable OM. Although water-soluble OM in the sediments is the primary interest of this study, the extraction method has the potential to bias towards the most labile pool of the sediment OM and can also extract a higher abundance of carbohydrates when compared to other solvents [68].

### 3.4. FTICR-MS Data Analysis

All FTICR-MS analyses were performed using R v4.0.0 [87], and all plots were generated using the ggplot2 package (v3.2.2) [88]. The R package “ftmsRanalysis” [89] was used to (1) remove peaks outside of a high confidence *m/z* range (200 *m/z*–900 *m/z*) and/or with a 13C isotopic signature; (2) calculate molecular formula properties (i.e., Kendrick defect, double-bond equivalent, modified aromaticity index, nominal oxidation state of carbon, standard Gibbs Free Energy of carbon oxidation); and (3) to determine to which chemical class a given metabolite belonged [45,46,48,50]. Using “ftmsRanalysis” [75] data outputs, we can obtain the central aspects of metabolomes investigated in this study, where elemental groups are categorized by the combination of elemental atoms present in each metabolite with molecular formula identified (e.g., CHO, CHON, CHOSP). The double-bond equivalent metric (DBE) describes the degree of chemical unsaturation of bonds in a particular metabolite [45,46], the modified aromaticity index (AI_Mod_) quantifies the degree of aromaticity (i.e., ring-like shape) of a metabolite [45–47], and NOSC indicates the energy required to oxidize different metabolomes [48]. High values of AI_Mod_ can denote the existence of either aromatic (AI_Mod_ > 0.5) or condensed aromatic structures (AI_Mod_≥ 0.67), and high DBE indicates more saturated compounds. NOSC is inversely correlated with the Gibbs free energy of carbon oxidation. Higher NOSC corresponds to metabolites that are more oxidized and thermodynamically favorable [15– 18,48,49]. Chemical class assignments for each metabolite were predicted using oxygen-to-carbon and hydrogen-to-carbon ratios (i.e., Van Krevelen classes [50]).

In order to evaluate bulk variation across sample types, a Mann–Whitney U test (*wilcox.test*) with a false discovery rate (FDR) *p*-value adjustment (*p.adjust*) was used to evaluate the divergence in molecular properties of all metabolites with molecular formulas assigned (46.08% of the total 95,681 peaks) present in either surface water or sediment. Differences in elemental group and chemical class relative abundances within samples between surface water and sediment were evaluated using the same approach. A principal component analysis (PCA; *prcomp*) was used to visualize differences between surface water and sediment metabolomes after a presence/absence transformation. A Euclidean distance matrix was obtained (*vegdist*, vegan package v2.5-6) and evaluated using a PERMANOVA (*adonis*, vegan package v2.5-6) in order to assess multivariate differences between sample types [90]. Inter-sample type variability was evaluated using the same Euclidean distance matrix in a beta-dispersion analysis (*betadisper*, vegan package v2.5-6); divergence in distance to centroid values was then evaluated using a Mann–Whitney U test [90].

To determine CONUS-scale patterns, sites were divided into eastern and western US based on their position relative to the location of the Mississippi River at St. Louis, Missouri. Replicates at each site were merged such that if a metabolite was observed in one replicate, it was considered present at the site. Given that FTICR-MS samples typically have less than 100% reproducibility [91,92], we considered a metabolite to be present in a sample if it was detected in any of the three replicates. This allowed us to maximize our detection of metabolites and has been previously employed [38]. Average molecular properties were then calculated, and elemental group/chemical class relative abundances were determined for each site/sample type combination based upon the metabolites present. This resulted in a single value for any given variable in surface water or sediment at a given site. Differences between these values across the East vs. West CONUS were then assessed using a Mann– Whitney U test with FDR correction. Maps were generated using ggplot to visualize spatial variance. All maps can be found in File S1.

### 3.5. Biochemical Transformation Analysis

We inferred biochemical transformations in sediment and surface water metabolomes as per Bailey et al. [62], Kaling et al. [69], Moritz et al. [70], Graham et al. [17,18], Garayburu-Caruso et al. [16], Danczak et al. [38], and Stegen et al. [15] to estimate the gain or loss of specific molecules (e.g., glucose, valine, glutamine). Briefly, pairwise mass differences were calculated between every peak in a sample and compared to a reference list of 1255 masses associated with commonly observed biochemical transformations (i.e., reactions of organic matter, Table S7). It is important to note that a molecular formula assignment is not necessary for this method as it allows for the incorporation of all detected metabolites. For mass differences matching to compounds in the reference list (within 1 ppm), we inferred the gain or loss of that compound via a biochemical transformation. For example, if a mass difference between two peaks corresponded to 71.0371, that would correlate to the loss or gain of the amino acid alanine, while a mass difference of 79.9662 would correspond to a loss or gain of a phosphate. Transformations were separated into 4 different groups based upon their labels: CHO-only, N-containing, S-containing, and P-containing. Differences in the relative abundance of transformations across samples were identified using a Mann–Whitney U test with FDR correction.

### 3.6. Data Availability

Original and expanded metadata, as well as surface water and sediment data used in this study, are publicly available on the Department of Energy data archive site ESS-DIVE [44,93]. All scripts used in this study are available on GitHub at https://github.com/danczakre/GlobalRiverMetabolomes.

## 4. Conclusions

We leveraged community science facilitated by the WHONDRS consortium to present the first ultrahigh-resolution analysis of global river corridor metabolomes of both surface water and sediment. Our data showed a strong divergence between surface water and sediment metabolomes, consistent with previous work within local systems. Surface water metabolomes were more rich and variable and contained more unsaturated and aromatic metabolites than sediment, possibly suggesting higher influence from terrestrial inputs or lower microbial processing. Further, surface water and sediment metabolomes had a consistent set of core biochemical transformations (CHO-only) but differed in N-, S-, and P-containing transformations that may be more influenced by nutrient limitations. Finally, we hypothesize the presence of systematic, spatially structured drivers influencing sediment metabolomes more strongly than surface water, as sediment changed along longitudinal patterns within the contiguous United States.

While there are many potential explanations for these patterns, the publicly available datasets being actively compiled by WHONDRS are well-suited for follow-on analyses to identify factors underlying metabolome variability. Given that the WHONDRS sampling campaign spanned 1^st^ to 9^th^ stream orders across multiple biomes (e.g., desert-like in the Columbia Plateau, subtropical in southern Florida, temperate forests in the Mid-Atlantic), outcomes of current and future data analyses and modeling efforts will enable transferable knowledge that can be applied throughout the world. To expand the breadth of questions that can be pursued with the data, WHONDRS is currently collecting information pertaining to mineralogy, geochemistry (e.g., anion and total N concentrations), microbiology (e.g., metagenomics, metatranscriptomics, flow cytometric cell counts), and various remote sensing data types (e.g., vegetation cover). Future questions might, for example, involve exploring spatial patterns of metabolomes across stream orders; correlating N-, S-, and P-containing transformations with land use, mineralogy, and vegetation; or investigating relationships between microbial activity and metabolome composition. We encourage the scientific community to explore WHONDRS datasets and combine them with additional data products to pursue novel scientific questions at local to global scales and to further engage with and pursue science that embodies the ICON principles.

## Supplementary Materials

The following are available online at www.mdpi.com/xxx/s1. Table S1: This table contains by-sample mean, median, and standard deviation for molecular characteristics (i.e., aromaticity index, H:C ratios, etc.; Sheet (1), the number of metabolites belonging to a given elemental group (i.e., CHON, CHO; Sheet (2), and the number of metabolites belonging to some compound class (i.e., %lignin-like, %protein-like, etc.; Sheet (3); Sheet (4), information about how each of the different measurements was calculated. Figure S1: Principal component analysis (PCA) performed on all peaks, regardless of formula assignment. Table S2: Loadings for PCA performed on all peaks. PC1 loadings on the x-axis are presented in Column A, while PC2 loadings on the y-axis are presented in Column B. Table S3: Loadings for PCA performed peaks with molecular formula assigned. PC1 loadings on the x-axis are presented in Column A, while PC2 loadings on the y-axis are presented in Column B. Table S4: This spreadsheet contains meta-data for each site from which samples were collected, including (but not limited to) latitude, longitude, stream order, and sampling date. Table S5: This table contains the results of the FDR-corrected, two-sided Mann–Whitney statistics performed to evaluate spatial variability across the contiguous United States of America. Table S6: Table of non-purgeable organic carbon (NPOC) concentrations for surface water and sediment-water extractions. Table S7: The file is the database of transformations used in the transformation analysis. The first column represents the transformation label, while the second column is the corresponding mass difference. There are two types of transformations listed in this file: (1) the gain or loss of the listed molecular formula (e.g., C1H1O1N1), with numeric values indicating the number of atoms associated with the element that precedes the numeric value; and (2) a substitution reaction denoted by an underscore (e.g., C1H1N1O_1). In the case of a substitution reaction, the underscore connects the element lost to the number of atoms lost. For example, C1H1N1O_1 indicates that a molecule gained C1H1N1 and lost one O atom. Some substitution reactions include multiple elements that are lost such that there are multiple underscores. In all cases, an underscore connects the element lost to the number of atoms lost. In all cases, atoms are gained if they are not followed immediately by an underscore. For example, C_1H_4O2 indicates loss of one C, loss of four H, and gain of two O. If no numeric value follows an element, it indicates that there is a gain of a single atom of that element (e.g., CH2 indicates one atom of C). File S1: A compressed file containing all of the maps generated during the spatial analysis of the contiguous United States of America.

## Author Contributions

V.A.G.-C., R.E.D., E.B.G., and J.C.S., conceptualized the study; V.A.G.-C., L.R., J.W., J.M.T., M.M., A.E.G., S.K., and S.F. carried out sample processing; C.T.R., R.K.C., and J.T. conducted the instrumental analyses; V.A.G.-C., R.E.D., J.C.S., and E.B.G. drafted the manuscript; and all authors contributed to the writing. All authors have read and agreed to the published version of the manuscript.

## Acknowledgments

The authors would like to thank the WHONDRS consortium.

## References

1. Battin, T.J.; Luyssaert, S.; Kaplan, L.A.; Aufdenkampe, A.K.; Richter, A.; Tranvik, L.J. The boundless carbon cycle. Nat. Geosci. 2009, 2, 598–600, doi:10.1038/ngeo618.

2. Cole, J.J.; Prairie, Y.T.; Caraco, N.F.; McDowell, W.H.; Tranvik, L.J.; Striegl, R.G.; Duarte, C.M.; Kortelainen, P.; Downing, J.A.; Middelburg, J.J.; et al. Plumbing the Global Carbon Cycle: Integrating Inland Waters into the Terrestrial Carbon Budget. Ecosystems 2007, 10, 172–185, doi:10.1007/s10021-006-9013-8.

3. Marín-Spiotta, E.; Gruley, K.E.; Crawford, J.; Atkinson, E.E.; Miesel, J.R.; Greene, S.; Cardona-Correa, C.; Spencer, R.G.M. Paradigm shifts in soil organic matter research affect interpretations of aquatic carbon cycling: Transcending disciplinary and ecosystem boundaries. Biogeochemistry 2014, 117, 279–297, doi:10.1007/s10533-013-9949-7.

4. Regnier, P.; Friedlingstein, P.; Ciais, P.; Mackenzie, F.T.; Gruber, N.; Janssens, I.A.; Laruelle, G.G.; Lauerwald, R.; Luyssaert, S.; Andersson, A.J.; et al. Anthropogenic perturbation of the carbon fluxes from land to ocean. Nat. Geosci. 2013, 6, 597–607, doi:10.1038/ngeo1830.

5. Hotchkiss, E.R.; Hall, R.O., Jr.; Sponseller, R.A.; Butman, D.; Klaminder, J.; Laudon, H.; Rosvall, M.; Karlsson, J. Sources of and processes controlling CO 2 emissions change with the size of streams and rivers. Nat. Geosci. 2015, 8, 696–699, doi:10.1038/ngeo2507.

6. Catalán, N.; Casas‐Ruiz, J.P.; Arce, M.I.; Abril, M.; Bravo, A.G.; Campo, R. del; Estévez, E.; Freixa, A.; Giménez‐Grau, P.; González‐Ferreras, A.M.; et al. Behind the Scenes: Mechanisms Regulating Climatic Patterns of Dissolved Organic Carbon Uptake in Headwater Streams. Glob. Biogeochem. Cycles 2018, 32, 1528–1541, doi:10.1029/2018GB005919.

7. Aufdenkampe, A.K.; Mayorga, E.; Raymond, P.A.; Melack, J.M.; Doney, S.C.; Alin, S.R.; Aalto, R.E.; Yoo, K. Riverine coupling of biogeochemical cycles between land, oceans, and atmosphere. Front. Ecol. Environ. 2011, 9, 53–60, doi:10.1890/100014.

8. Raymond, P.A.; Hartmann, J.; Lauerwald, R.; Sobek, S.; McDonald, C.; Hoover, M.; Butman, D.; Striegl, R.; Mayorga, E.; Humborg, C.; et al. Global carbon dioxide emissions from inland waters. Nature 2013, 503, 355–359, doi:10.1038/nature12760.

9. Moody, C.S.; Worrall, F.; Evans, C.D.; Jones, T.G. The rate of loss of dissolved organic carbon (DOC) through a catchment. J. Hydrol. 2013, 492, 139–150, doi:10.1016/j.jhydrol.2013.03.016.

10. Cory, R.M.; Ward, C.P.; Crump, B.C.; Kling, G.W. Sunlight controls water column processing of carbon in arctic fresh waters. Science 2014, 345, 925, doi:10.1126/science.1253119.

11. Newbold, J.D.; Bott, T.L.; Kaplan, L.A.; Dow, C.L.; Jackson, J.K.; Aufdenkampe, A.K.; Martin, L.A.; Horn, D.J.V.; Long, A.A. Uptake of nutrients and organic C in streams in New York City drinking-water-supply watersheds. Freshw. Sci. 2006, 25, 998–1017, doi:10.1899/0887-3593(2006)025 [0998:UONAOC]2.0.CO;2.

12. Bernhardt, E.S.; McDowell, W.H. Twenty years apart: Comparisons of DOM uptake during leaf leachate releases to Hubbard Brook Valley streams in 1979 versus 2000. J. Geophys. Res. Biogeosci. 2008, 113, doi:10.1029/2007JG000618.

13. Koehler, B.; Wachenfeldt, E. von; Kothawala, D.; Tranvik, L.J. Reactivity continuum of dissolved organic carbon decomposition in lake water. J. Geophys. Res. Biogeosci. 2012, 117, doi:10.1029/2011JG001793.

14. Berggren, M.; Giorgio, P.A. del Distinct patterns of microbial metabolism associated to riverine dissolved organic carbon of different source and quality. J. Geophys. Res. Biogeosci. 2015, 120, 989–999, doi:10.1002/2015JG002963.

15. Stegen, J.C.; Johnson, T.; Fredrickson, J.K.; Wilkins, M.J.; Konopka, A.E.; Nelson, W.C.; Arntzen, E.V.; Chrisler, W.B.; Chu, R.K.; Fansler, S.J.; et al. Influences of organic carbon speciation on hyporheic corridor biogeochemistry and microbial ecology. Nat. Commun. 2018, 9, 1034, doi:10.1038/s41467-018-02922-9.

16. Garayburu-Caruso, V.A.; Stegen, J.C.; Song, H.-S.; Renteria, L.; Wells, J.; Garcia, W.; Resch, C.T.; Goldman, A.E.; Chu, R.K.; Toyoda, J.; et al. Carbon Limitation Leads to Thermodynamic Regulation of Aerobic Metabolism. Environ. Sci. Technol. Lett. 2020, 7, 517–524, doi:10.1021/acs.estlett.0c00258.

17. Graham, E.B.; Crump, A.R.; Kennedy, D.W.; Arntzen, E.; Fansler, S.; Purvine, S.O.; Nicora, C.D.; Nelson, W.; Tfaily, M.M.; Stegen, J.C. Multi’omics comparison reveals metabolome biochemistry, not microbiome composition or gene expression, corresponds to elevated biogeochemical function in the hyporheic zone. Sci. Total Environ. 2018, 642, 742–753, doi:10.1016/j.scitotenv.2018.05.256.

18. Graham, E.B.; Tfaily, M.M.; Crump, A.R.; Goldman, A.E.; Bramer, L.M.; Arntzen, E.; Romero, E.; Resch, C.T.; Kennedy, D.W.; Stegen, J.C. Carbon Inputs From Riparian Vegetation Limit Oxidation of Physically Bound Organic Carbon Via Biochemical and Thermodynamic Processes. J. Geophys. Res. Biogeosci. 2017, 122, 3188–3205, doi:10.1002/2017JG003967.

19. Bundy, J.G.; Davey, M.P.; Viant, M.R. Environmental metabolomics: A critical review and future perspectives. Metabolomics 2008, 5, 3, doi:10.1007/s11306-008-0152-0.

20. Rue, G.P.; Rock, N.D.; Gabor, R.S.; Pitlick, J.; Tfaily, M.; McKnight, D.M. Concentration-discharge relationships during an extreme event: Contrasting behavior of solutes and changes to chemical quality of dissolved organic material in the Boulder Creek Watershed during the September 2013 flood. Water Resour. Res. 2017, 53, 5276–5297, doi:10.1002/2016WR019708.

21. Wilson, R.M.; Tfaily, M.M. Advanced Molecular Techniques Provide New Rigorous Tools for Characterizing Organic Matter Quality in Complex Systems. J. Geophys. Res. Biogeosci. 2018, 123, 1790–1795, doi:10.1029/2018JG004525.

22. Li, L.; He, Z.L.; Tfaily, M.M.; Inglett, P.; Stoffella, P.J. Spatial-temporal variations of dissolved organic nitrogen molecular composition in agricultural runoff water. Water Res. 2018, 137, 375–383, doi:10.1016/j.watres.2018.01.035.

23. Walker, L.R.; Tfaily, M.M.; Shaw, J.B.; Hess, N.J.; Paša-Tolić, L.; Koppenaal, D.W. Unambiguous identification and discovery of bacterial siderophores by direct injection 21 Tesla Fourier transform ion cyclotron resonance mass spectrometry. Metallomics 2017, 9, 82–92, doi:10.1039/C6MT00201C.

24. Hodgkins, S.B.; Tfaily, M.M.; McCalley, C.K.; Logan, T.A.; Crill, P.M.; Saleska, S.R.; Rich, V.I.; Chanton, J.P. Changes in peat chemistry associated with permafrost thaw increase greenhouse gas production. Proc. Natl. Acad. Sci. USA 2014, 111, 5819–5824, doi:10.1073/pnas.1314641111.

25. Jones, O.A.H.; Lear, G.; Welji, A.M.; Collins, G.; Quince, C. Community Metabolomics in Environmental Microbiology. In Microbial Metabolomics: Applications in Clinical, Environmental, and Industrial Microbiology; Beale, D.J., Kouremenos, K.A., Palombo, E.A., Eds.; Springer International Publishing: Cham, Switzerland, 2016; pp. 199–224. ISBN 978-3-319-46326-1.

26. Jones, O.A.H.; Sdepanian, S.; Lofts, S.; Svendsen, C.; Spurgeon, D.J.; Maguire, M.L.; Griffin, J.L. Metabolomic analysis of soil communities can be used for pollution assessment. Environ. Toxicol. Chem. 2014, 33, 61–64, doi:10.1002/etc.2418.

27. Beale, D.J.; Crosswell, J.; Karpe, A.V.; Metcalfe, S.S.; Morrison, P.D.; Staley, C.; Ahmed, W.; Sadowsky, M.J.; Palombo, E.A.; Steven, A.D.L. Seasonal metabolic analysis of marine sediments collected from Moreton Bay in South East Queensland, Australia, using a multi-omics-based approach. Sci. Total Environ. 2018, 631–632, 1328–1341, doi:10.1016/j.scitotenv.2018.03.106.

28. Beale, D.J.; Crosswell, J.; Karpe, A.V.; Ahmed, W.; Williams, M.; Morrison, P.D.; Metcalfe, S.; Staley, C.; Sadowsky, M.J.; Palombo, E.A.; et al. A multi-omics based ecological analysis of coastal marine sediments from Gladstone, in Australia’s Central Queensland, and Heron Island, a nearby fringing platform reef. Sci. Total Environ. 2017, 609, 842–853, doi:10.1016/j.scitotenv.2017.07.184.

29. Shah, R.M.; Crosswell, J.; Metcalfe, S.S.; Carlin, G.; Morrison, P.D.; Karpe, A.V.; Palombo, E.A.; Steven, A.D.L.; Beale, D.J. Influence of Human Activities on Broad-Scale Estuarine-Marine Habitats Using Omics-Based Approaches Applied to Marine Sediments. Microorganisms 2019, 7, doi:10.3390/microorganisms7100419.

30. Kimes, N.E.; Callaghan, A.V.; Aktas, D.F.; Smith, W.L.; Sunner, J.; Golding, B.; Drozdowska, M.; Hazen, T.C.; Suflita, J.M.; Morris, P.J. Metagenomic analysis and metabolite profiling of deep-sea sediments from the Gulf of Mexico following the Deepwater Horizon oil spill. Front. Microbiol. 2013, 4, 50, doi:10.3389/fmicb.2013.00050.

31. Lam, B.; Baer, A.; Alaee, M.; Lefebvre, B.; Moser, A.; Williams, A.; Simpson, A.J. Major Structural Components in Freshwater Dissolved Organic Matter. Environ. Sci. Technol. 2007, 41, 8240–8247, doi:10.1021/es0713072.

32. Jaffé, R.; McKnight, D.; Maie, N.; Cory, R.; McDowell, W.H.; Campbell, J.L. Spatial and temporal variations in DOM composition in ecosystems: The importance of long-term monitoring of optical properties. J. Geophys. Res. Biogeosci. 2008, 113, doi:10.1029/2008JG000683.

33. Gonsior, M.; Peake, B.M.; Cooper, W.T.; Podgorski, D.; D’Andrilli, J.; Cooper, W.J. Photochemically induced changes in dissolved organic matter identified by ultrahigh resolution fourier transform ion cyclotron resonance mass spectrometry. Environ. Sci. Technol. 2009, 43, 698–703, doi:10.1021/es8022804.

34. Lu, Y.; Li, X.; Mesfioui, R.; Bauer, J.E.; Chambers, R.M.; Canuel, E.A.; Hatcher, P.G. Use of ESI-FTICR-MS to Characterize Dissolved Organic Matter in Headwater Streams Draining Forest-Dominated and Pasture-Dominated Watersheds. PLoS ONE 2015, 10, e0145639, doi:10.1371/journal.pone.0145639.

35. Stegen, J.C.; Goldman, A.E. WHONDRS: A Community Resource for Studying Dynamic River Corridors. mSystems 2018, 3, doi:10.1128/mSystems.00151-18.

36. Jaffé, R.; Yamashita, Y.; Maie, N.; Cooper, W.T.; Dittmar, T.; Dodds, W.K.; Jones, J.B.; Myoshi, T.; Ortiz-Zayas, J.R.; Podgorski, D.C.; et al. Dissolved Organic Matter in Headwater Streams: Compositional Variability across Climatic Regions of North America. Geochim. Cosmochim. Acta 2012, 94, 95–108, doi:10.1016/j.gca.2012.06.031.

37. Wagner, S.; Riedel, T.; Niggemann, J.; Vähätalo, A.V.; Dittmar, T.; Jaffé, R. Linking the Molecular Signature of Heteroatomic Dissolved Organic Matter to Watershed Characteristics in World Rivers. Environ. Sci. Technol. 2015, 49, 13798–13806, doi:10.1021/acs.est.5b00525.

38. Danczak, R.E.; Goldman, A.E.; Chu, R.K.; Toyoda, J.G.; Garayburu-Caruso, V.A.; Tolić, N.; Graham, E.B.; Morad, J.W.; Renteria, L.; Wells, J.R.; et al. Ecological theory applied to environmental metabolomes reveals compositional divergence despite conserved molecular properties. bioRxiv 2020, doi:10.1101/2020.02.12.946459.

39. Steefel, C.I.; Appelo, C.A.J.; Arora, B.; Jacques, D.; Kalbacher, T.; Kolditz, O.; Lagneau, V.; Lichtner, P.C.; Mayer, K.U.; Meeussen, J.C.L.; et al. Reactive transport codes for subsurface environmental simulation. Comput. Geosci. 2015, 19, 445–478, doi:10.1007/s10596-014-9443-x.

40. Song, H.-S.; Stegen, J.C.; Graham, E.B.; Lee, J.-Y.; Garayburu-Caruso, V.A.; Nelson, W.C.; Chen, X.; Moulton, J.D.; Scheibe, T.D. Representing Organic Matter Thermodynamics in Biogeochemical Reactions via Substrate-Explicit Modeling. bioRxiv 2020, doi:10.1101/2020.02.27.968669.

41. Stegen, J.; Brodie, E.; Wrighton, K.; Bayer, P.; Lesmes, D.; Emani, S.; Moerman, J. Open Watershed Science by Design: Leveraging Distributed Research Networks to Understand Watershed Systems: Workshop Report, DOE/SC-0200; USDOE Office of Science (SC): USA, 2019, doesbr.org/openwatersheds/.

42. Uhlmann, E.L.; Ebersole, C.R.; Chartier, C.R.; Errington, T.M.; Kidwell, M.C.; Lai, C.K.; McCarthy, R.J.; Riegelman, A.; Silberzahn, R.; Nosek, B.A. Scientific Utopia III: Crowdsourcing Science: Perspect. Psychol. Sci. 2019, doi:10.1177/1745691619850561.

43. Wilkinson, M.D.; Dumontier, M.; Aalbersberg, I.J.J.; Appleton, G.; Axton, M.; Baak, A.; Blomberg, N.; Boiten, J.-W.; da Silva Santos, L.B.; Bourne, P.E.; et al. The FAIR Guiding Principles for scientific data management and stewardship. Sci. Data 2016, 3, 160018, doi:10.1038/sdata.2016.18.

44. Toyoda, J.G.; Goldman, A.E.; Chu, R.K.; Danczak, R.E.; Daly, R.A.; Garayburu-Caruso, V.A.; Graham, E.B.; Lin, X.; Moran, J.J.; Ren, H. WHONDRS Summer 2019 Sampling Campaign: Global River Corridor Surface Water FTICR-MS and Stable Isotopes; Environmental System Science Data Infrastructure for a Virtual Ecosystem, Worldwide Hydrobiogeochemistry Observation Network for Dynamic River Systems (WHONDRS): 2020, doi:10.15485/1603775

45. Koch, B.P.; Dittmar, T. From mass to structure: An aromaticity index for high-resolution mass data of natural organic matter. Rapid Commun. Mass Spectrom. 2006, 20, 926–932, doi:10.1002/rcm.2386.

46. Koch, B.P.; Dittmar, T. From mass to structure: An aromaticity index for high-resolution mass data of natural organic matter. Rapid Commun. Mass Spectrom. 2016, 30, 250–250, doi:10.1002/rcm.7433.

47. Willoughby, A.S.; Wozniak, A.S.; Hatcher, P.G. A molecular-level approach for characterizing water-insoluble components of ambient organic aerosol particulates using ultrahigh-resolution mass spectrometry. Atmos. Chem. Phys. 2014, 14, 10299–10314, doi:10.5194/acp-14-10299-2014.

48. LaRowe, D.E.; Van Cappellen, P. Degradation of natural organic matter: A thermodynamic analysis. Geochim. Cosmochim. Acta 2011, 75, 2030–2042, doi:10.1016/j.gca.2011.01.020.

49. Boye, K.; Noël, V.; Tfaily, M.M.; Bone, S.E.; Williams, K.H.; Bargar, J.R.; Fendorf, S. Thermodynamically controlled preservation of organic carbon in floodplains. Nat. Geosci. 2017, 10, 415–419, doi:10.1038/ngeo2940.

50. Kim, S.; Kramer, R.W.; Hatcher, P.G. Graphical Method for Analysis of Ultrahigh-Resolution Broadband Mass Spectra of Natural Organic Matter, the Van Krevelen Diagram. Anal. Chem. 2003, 75, 5336–5344, doi:10.1021/ac034415p.

51. Hedges, J.I.; Mann, D.C. The lignin geochemistry of marine sediments from the southern Washington coast. Geochim. Cosmochim. Acta 1979, 43, 1809–1818, doi:10.1016/0016-7037(79)90029-2.

52. Stegen, J.C.; Fredrickson, J.K.; Wilkins, M.J.; Konopka, A.E.; Nelson, W.C.; Arntzen, E.V.; Chrisler, W.B.; Chu, R.K.; Danczak, R.E.; Fansler, S.J.; et al. Groundwater–surface water mixing shifts ecological assembly processes and stimulates organic carbon turnover. Nat. Commun. 2016, 7, 11237, doi:10.1038/ncomms11237.

53. Pracht, L.E.; Tfaily, M.M.; Ardissono, R.J.; Neumann, R.B. Molecular characterization of organic matter mobilized from Bangladeshi aquifer sediment: Tracking carbon compositional change during microbial utilization. Biogeosciences 2018, 15, 1733–1747, doi:10.5194/bg-15-1733-2018.

54. Valle, J.; Harir, M.; Gonsior, M.; Enrich-Prast, A.; Schmitt-Kopplin, P.; Bastviken, D.; Hertkorn, N. Molecular differences between water column and sediment pore water SPE-DOM in ten Swedish boreal lakes. Water Res. 2020, 170, 115320, doi:10.1016/j.watres.2019.115320.

55. Boulton, A.J.; Findlay, S.; Marmonier, P.; Stanley, E.H.; Valett, H.M. The functional significance of the hyporheic zone in streams and rivers. Annu. Rev. Ecol. Syst. 1998, 29, 59–81, doi:10.1146/annurev.ecolsys.29.1.59.

56. Boano, F.; Harvey, J.W.; Marion, A.; Packman, A.I.; Revelli, R.; Ridolfi, L.; Wörman, A. Hyporheic flow and transport processes: Mechanisms, models, and biogeochemical implications. Rev. Geophys. 2014, 52, 603–679, doi:10.1002/2012RG000417.

57. Harvey, J.; Gooseff, M. River corridor science: Hydrologic exchange and ecological consequences from bedforms to basins. Water Resour. Res. 2015, 51, 6893–6922, doi:10.1002/2015WR017617.

58. Gomez-Velez, J.D.; Harvey, J.W.; Cardenas, M.B.; Kiel, B. Denitrification in the Mississippi River network controlled by flow through river bedforms. Nat. Geosci. 2015, 8, 941–945, doi:10.1038/ngeo2567.

59. Knapp, J.L.A.; González‐Pinzón, R.; Drummond, J.D.; Larsen, L.G.; Cirpka, O.A.; Harvey, J.W. Tracer-based characterization of hyporheic exchange and benthic biolayers in streams. Water Resour. Res. 2017, 53, 1575–1594, doi:10.1002/2016WR019393.

60. Fischer, H.; Kloep, F.; Wilzcek, S.; Pusch, M.T. A river’s liver–microbial processes within the hyporheic zone of a large lowland river. Biogeochemistry 2005, 76, 349–371.

61. Battin, T.J.; Besemer, K.; Bengtsson, M.M.; Romani, A.M.; Packmann, A.I. The ecology and biogeochemistry of stream biofilms. Nat. Rev. Microbiol. 2016, 14, 251.

62. Bailey, V.L.; Smith, A.P.; Tfaily, M.; Fansler, S.J.; Bond-Lamberty, B. Differences in soluble organic carbon chemistry in pore waters sampled from different pore size domains. Soil Biol. Biochem. 2017, 107, 133–143, doi:10.1016/j.soilbio.2016.11.025.

63. Cory, R.M.; Kling, G.W. Interactions between sunlight and microorganisms influence dissolved organic matter degradation along the aquatic continuum. Limnol. Oceanogr. Lett. 2018, 3, 102–116, doi:10.1002/lol2.10060.

64. Bao, H.; Niggemann, J.; Huang, D.; Dittmar, T.; Kao, S.-J. Different Responses of Dissolved Black Carbon and Dissolved Lignin to Seasonal Hydrological Changes and an Extreme Rain Event. J. Geophys. Res. Biogeosciences 2019, 124, 479–493, doi:10.1029/2018JG004822.

65. Fellman, J.B.; Hood, E.; Edwards, R.T.; D’Amore, D.V. Changes in the concentration, biodegradability, and fluorescent properties of dissolved organic matter during stormflows in coastal temperate watersheds. J. Geophys. Res. Biogeosci. 2009, 114, doi:10.1029/2008JG000790.

66. Hood, E.; Gooseff, M.N.; Johnson, S.L. Changes in the character of stream water dissolved organic carbon during flushing in three small watersheds, Oregon. J. Geophys. Res. Biogeosci. 2006, 111, doi:10.1029/2005JG000082.

67. Ward, N.D.; Richey, J.E.; Keil, R.G. Temporal variation in river nutrient and dissolved lignin phenol concentrations and the impact of storm events on nutrient loading to Hood Canal, Washington, USA. Biogeochemistry 2012, 111, 629–645, doi:10.1007/s10533-012-9700-9.

68. Tfaily, M.M.; Chu, R.K.; Toyoda, J.; Tolić, N.; Robinson, E.W.; Paša-Tolić, L.; Hess, N.J. Sequential extraction protocol for organic matter from soils and sediments using high resolution mass spectrometry. Anal. Chim. Acta 2017, 972, 54–61, doi:10.1016/j.aca.2017.03.031.

69. Kaling, M.; Schmidt, A.; Moritz, F.; Rosenkranz, M.; Witting, M.; Kasper, K.; Janz, D.; Schmitt-Kopplin, P.; Schnitzler, J.-P.; Polle, A. Mycorrhiza-Triggered Transcriptomic and Metabolomic Networks Impinge on Herbivore Fitness. Plant Physiol. 2018, 176, 2639–2656, doi:10.1104/pp.17.01810.

70. Moritz, F.; Kaling, M.; Schnitzler, J.-P.; Schmitt‐Kopplin, P. Characterization of poplar metabotypes via mass difference enrichment analysis. Plant Cell Environ. 2017, 40, 1057–1073, doi:10.1111/pce.12878.

71. Craine, J.M.; Morrow, C.; Fierer, N. Microbial Nitrogen Limitation Increases Decomposition. Ecology 2007, 88, 2105–2113, doi:10.1890/06-1847.1.

72. Moorhead, D.L.; Sinsabaugh, R.L. A Theoretical Model of Litter Decay and Microbial Interaction. Ecol. Monogr. 2006, 76, 151–174, doi:10.1890/0012-9615(2006)076[0151:ATMOLD]2.0.CO;2.

73. Zhao, F.J.; Wu, J.; McGrath, S.P. Chapter 12—Soil Organic Sulphur and its Turnover. In Humic Substances in Terrestrial Ecosystems; Piccolo, A., Ed.; Elsevier Science B.V.: Amsterdam, The Netherlands, 1996; pp. 467–506. ISBN 978-0-444-81516-3.

74. Reddy, K.R.; Kadlec, R.H.; Flaig, E.; Gale, P.M. Phosphorus Retention in Streams and Wetlands: A Review. Crit. Rev. Environ. Sci. Technol. 1999, 29, 83–146, doi:10.1080/10643389991259182.

75. Freixa, A.; Ejarque, E.; Crognale, S.; Amalfitano, S.; Fazi, S.; Butturini, A.; Romaní, A.M. Sediment microbial communities rely on different dissolved organic matter sources along a Mediterranean river continuum. Limnol. Oceanogr. 2016, 61, 1389–1405, doi:10.1002/lno.10308.

76. Vazquez, E.; Amalfitano, S.; Fazi, S.; Butturini, A. Dissolved organic matter composition in a fragmented Mediterranean fluvial system under severe drought conditions. Biogeochemistry 2011, 102, 59–72, doi:10.1007/s10533-010-9421-x.

77. Butturini, A.; Guarch, A.; Romaní, A.M.; Freixa, A.; Amalfitano, S.; Fazi, S.; Ejarque, E. Hydrological conditions control in situ DOM retention and release along a Mediterranean river. Water Res. 2016, 99, 33–45, doi:10.1016/j.watres.2016.04.036.

78. Ejarque, E.; Freixa, A.; Vazquez, E.; Guarch, A.; Amalfitano, S.; Fazi, S.; Romaní, A.M.; Butturini, A. Quality and reactivity of dissolved organic matter in a Mediterranean river across hydrological and spatial gradients. Sci. Total Environ. 2017, 599–600, 1802–1812, doi:10.1016/j.scitotenv.2017.05.113.

79. Robinson, N.P.; Allred, B.W.; Jones, M.O.; Moreno, A.; Kimball, J.S.; Naugle, D.E.; Erickson, T.A.; Richardson, A.D. A dynamic Landsat derived normalized difference vegetation index (NDVI) product for the conterminous United States. Remote Sens. 2017, 9, 863.

80. Sayre, R. (Ed.) A New Map of Standardized Terrestrial Ecosystems of the Conterminous United States; U.S. Department of the Interior, U.S. Geological Survey: Reston, VA, USA, 2009; ISBN 978-1-4113-2432-9.

81. Fierer, N.; Ladau, J.; Clemente, J.C.; Leff, J.W.; Owens, S.M.; Pollard, K.S.; Knight, R.; Gilbert, J.A.; McCulley, R.L. Reconstructing the Microbial Diversity and Function of Pre-Agricultural Tallgrass Prairie Soils in the United States. Science 2013, 342, 621–624, doi:10.1126/science.1243768.

82. Horton, R.E. Erosional Development of Streams and Their Drainage Basins; Hydrophysical Approach to Quantitative Morphology. GSA Bull. 1945, 56, 275–370, doi:10.1130/0016-7606(1945)56[275:EDOSAT]2.0.CO;2.

83. Strahler, A.N. Dimensional Analysis Applied to Fluvially Eroded Landforms. GSA Bull. 1958, 69, 279–300, doi:10.1130/0016-7606(1958)69[279:DAATFE]2.0.CO;2.

84. Jensen, B. AOS Protocol and Procedure: Sediment Chemistry Sampling in Wadeable Streams (NEON.DOC.001193). Available online: http://data.neonscience.org/documents (accessed June 2019.

85. Dittmar, T.; Koch, B.; Hertkorn, N.; Kattner, G. A simple and efficient method for the solid-phase extraction of dissolved organic matter (SPE-DOM) from seawater: SPE-DOM from seawater. Limnol. Oceanogr. Methods 2008, 6, 230–235, doi:10.4319/lom.2008.6.230.

86. Tolić, N.; Liu, Y.; Liyu, A.; Shen, Y.; Tfaily, M.M.; Kujawinski, E.B.; Longnecker, K.; Kuo, L.-J.; Robinson, E.W.; Paša-Tolić, L.; et al. Formularity: Software for Automated Formula Assignment of Natural and Other Organic Matter from Ultrahigh-Resolution Mass Spectra. Anal. Chem. 2017, 89, 12659–12665, doi:10.1021/acs.analchem.7b03318.

87. R Core Team. R: A Language and Environment for Statistical Computing; R Foundation for Statistical Computing: Vienna, Austria, 2020.

88. Wickham, H. ggplot2: Elegant Graphics for Data Analysis; Springer: New York, NY, USA, 2016.

89. Bramer, L.M.; White, A.M.; Stratton, K.G.; Thompson, A.M.; Claborne, D.; Hofmockel, K.; McCue, L.A. ftmsRanalysis: An R package for exploratory data analysis and interactive visualization of FT-MS data. PLoS Comput. Biol. 2020, 16, e1007654, doi:10.1371/journal.pcbi.1007654.

90. Oksanen, J.; Blanchet, F.G.; Friendly, M.; Kindt, R.; Legendre, P.; McGlinn, D.; Minchin, P.R.; O’hara, R.; Simpson, G.L.; Solymos, P. vegan: Community Ecology Package. R package version 2.5-6. Vienna R Found. Stat. Comput. Sch. 2019.

91. Hawkes, J.A.; D’Andrilli, J.; Agar, J.N.; Barrow, M.P.; Berg, S.M.; Catalán, N.; Chen, H.; Chu, R.K.; Cole, R.B.; Dittmar, T.; et al. An international laboratory comparison of dissolved organic matter composition by high resolution mass spectrometry: Are we getting the same answer? Limnol. Oceanogr. Methods 2020, 18, 235–258, doi:10.1002/lom3.10364.

92. He, C.; Zhang, Y.; Li, Y.; Zhuo, X.; Li, Y.; Zhang, C.; Shi, Q. In-House Standard Method for Molecular Characterization of Dissolved Organic Matter by FT-ICR Mass Spectrometry. ACS Omega 2020, 5, 11730–11736, doi:10.1021/acsomega.0c01055.

93. Goldman, A.E.; Chu, R.K.; Danczak, R.E.; Daly, R.A.; Fansler, S.; Garayburu-Caruso, V.A.; Graham, E.B.; McCall, M.L.; Ren, H.; Renteria, L. WHONDRS Summer 2019 Sampling Campaign: Global River Corridor Sediment FTICR-MS, NPOC, and Aerobic Respiration; Environmental System Science Data Infrastructure for a Virtual Ecosystem, Worldwide Hydrobiogeochemistry Observation Network for Dynamic River Systems (WHONDRS), 2020, doi:10.15485/1729719

