## Supplementary figures and images for "Using Community Science to Reveal the Global Chemogeography of River Metabolomes"

### Supplementary Figure 1

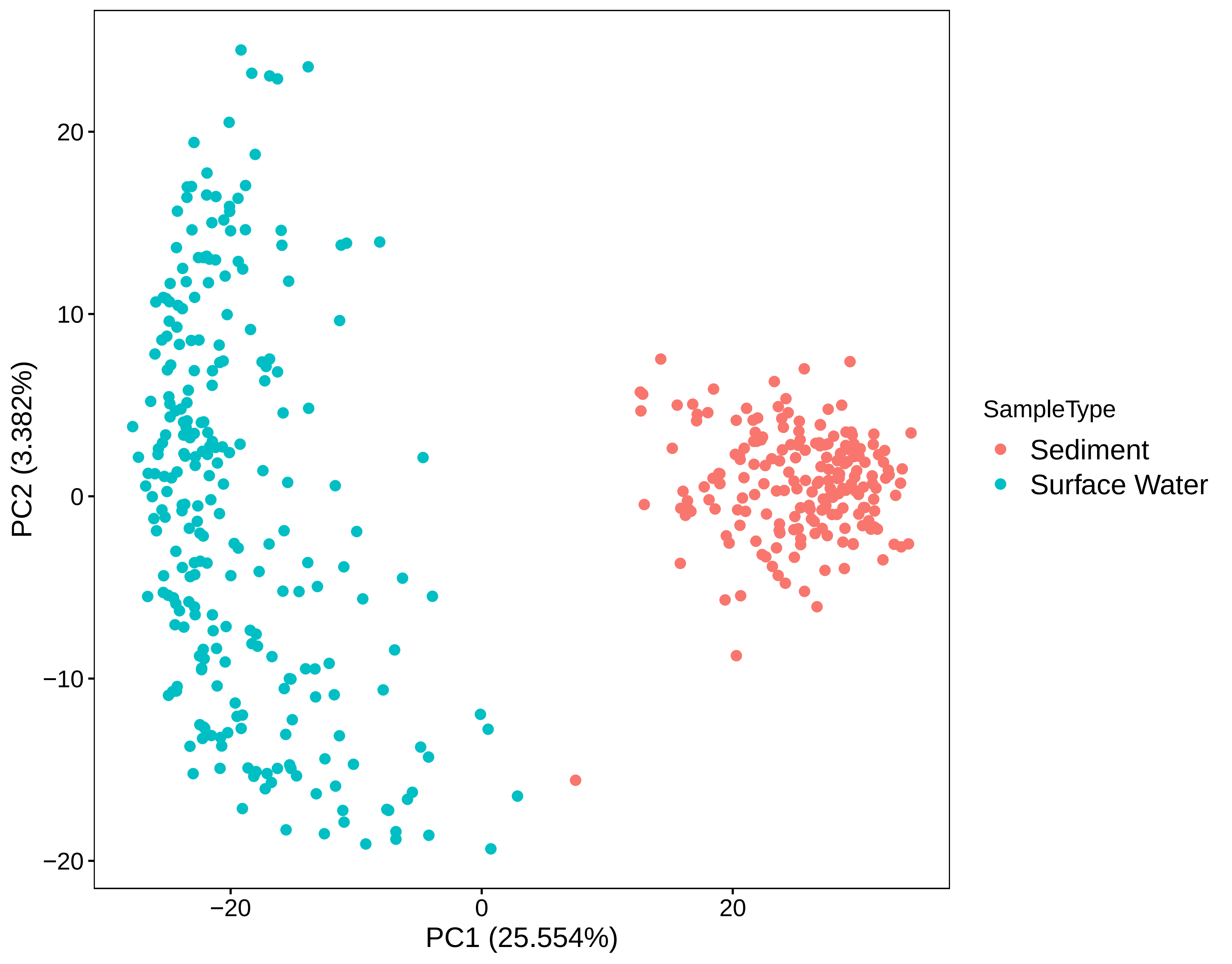

### Supplementary File 1

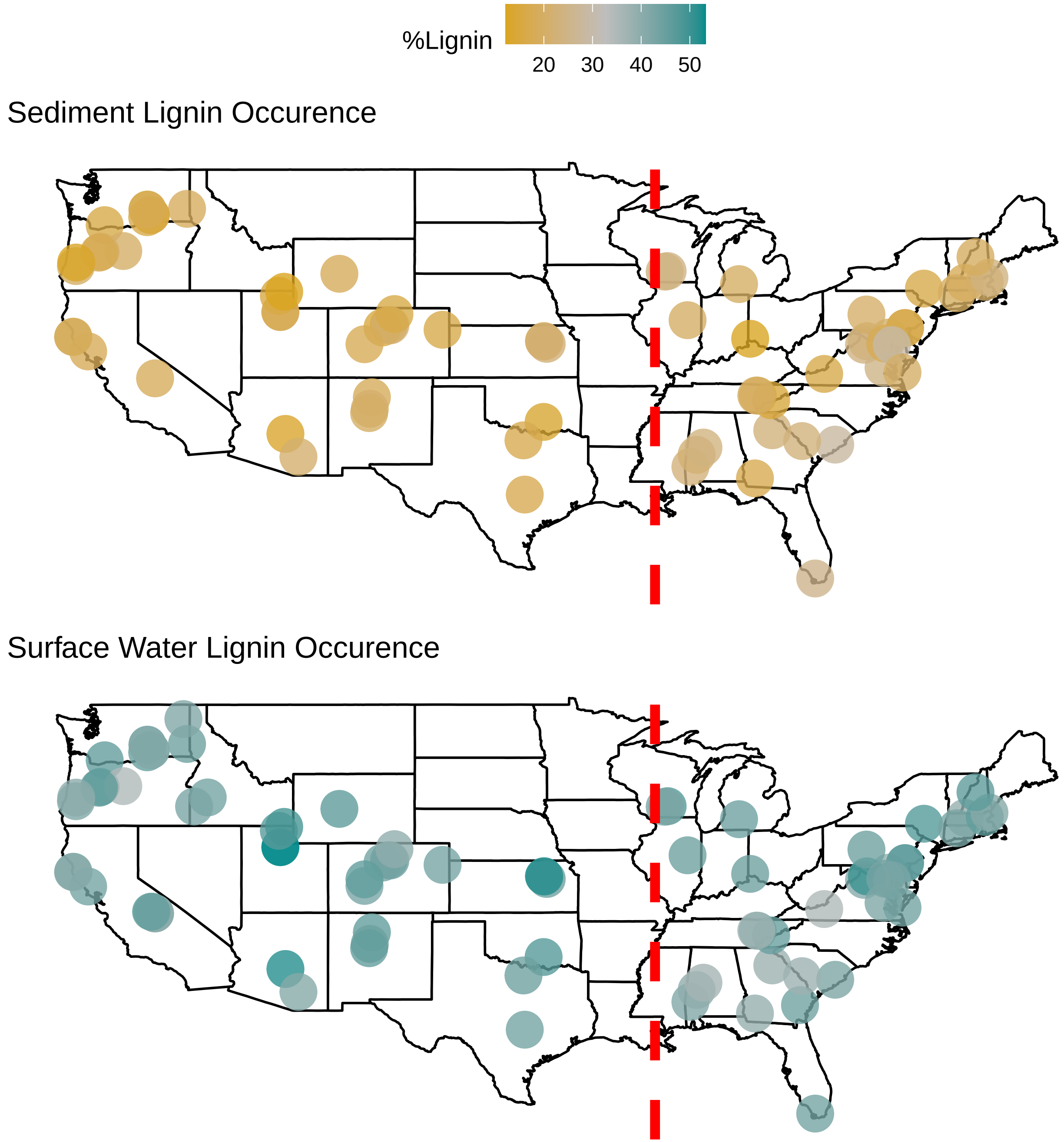

### Supplementary File 1

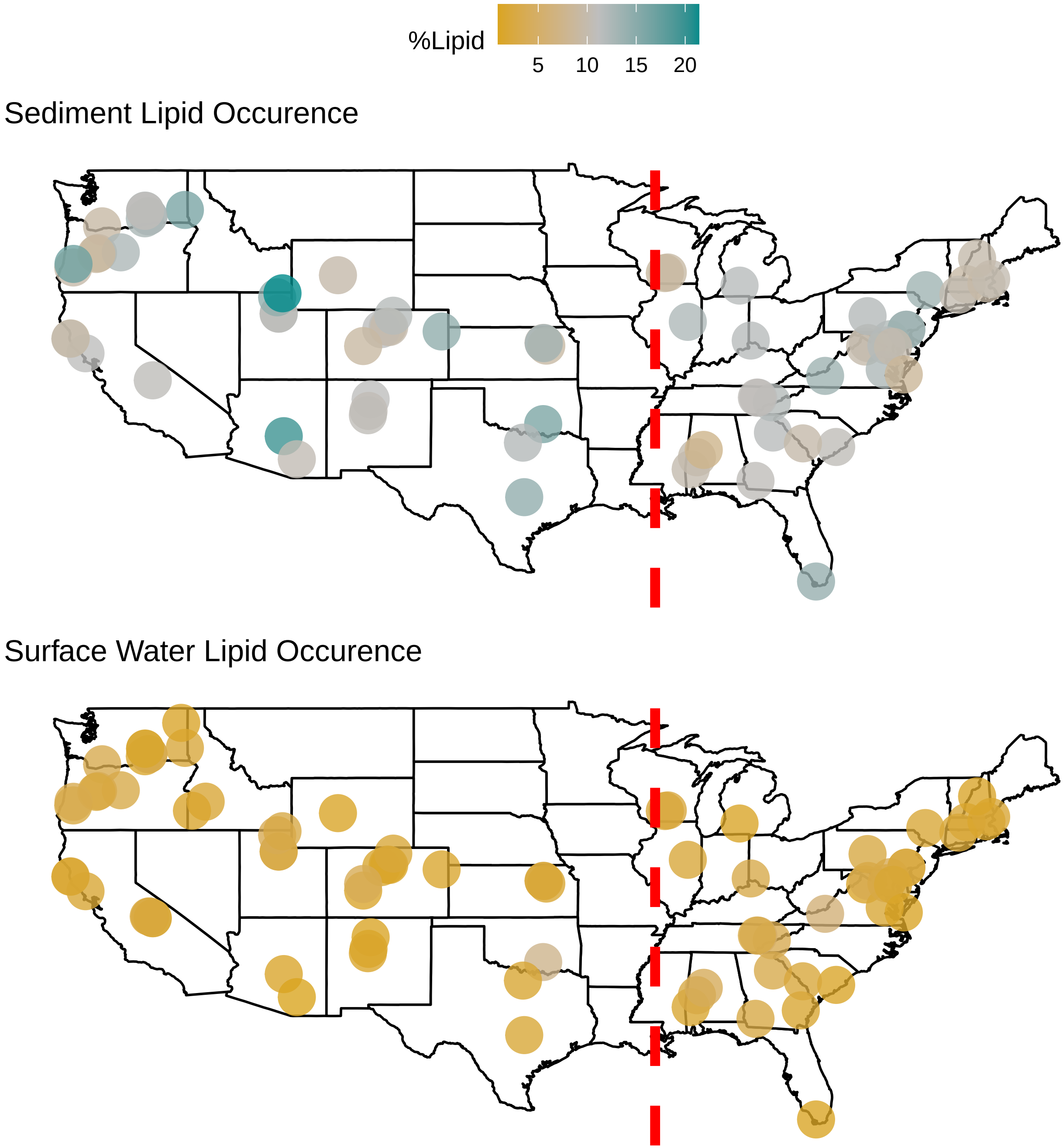

### Supplementary File 1

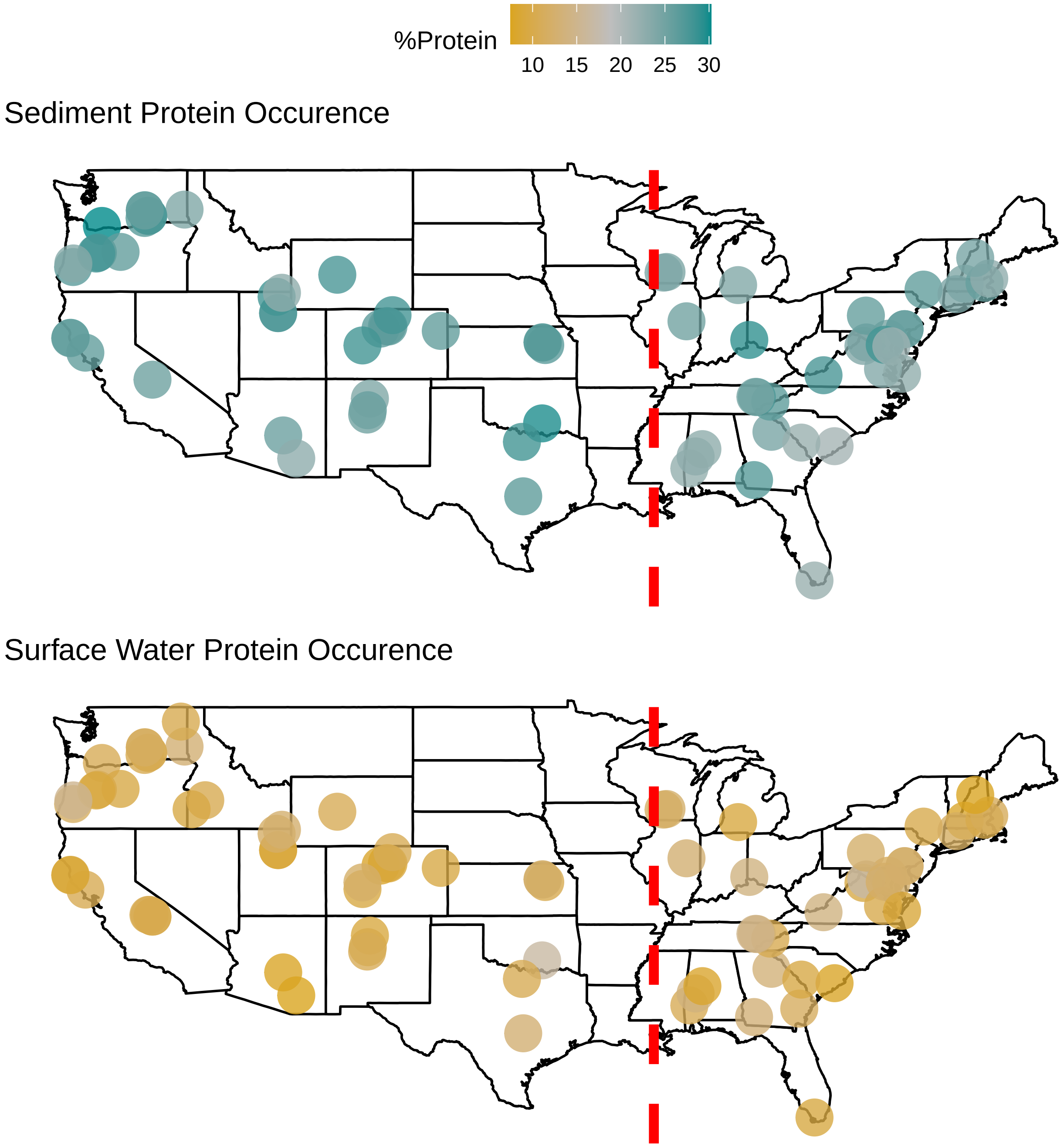

### Supplementary File 1

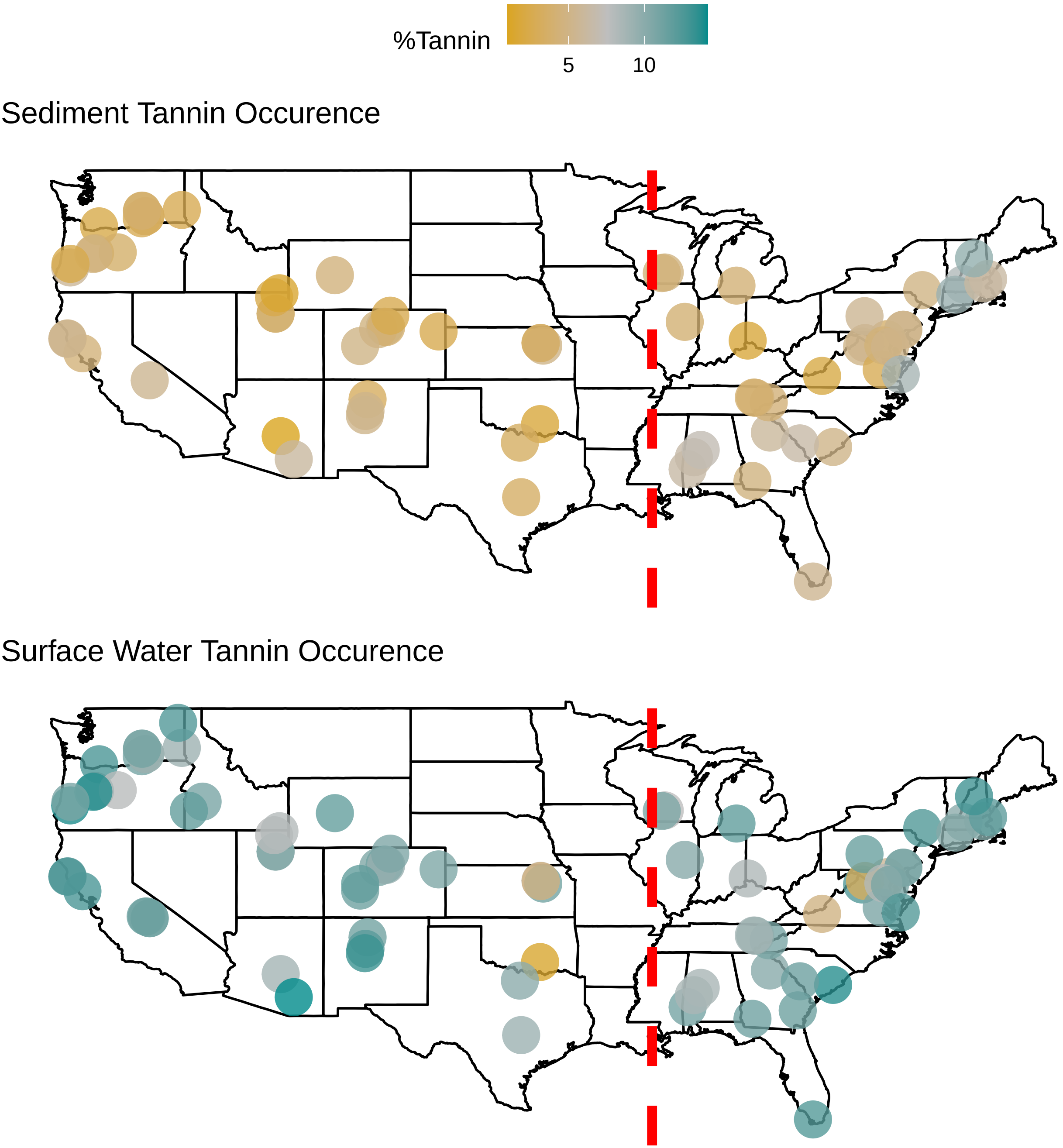

### Supplementary File 1

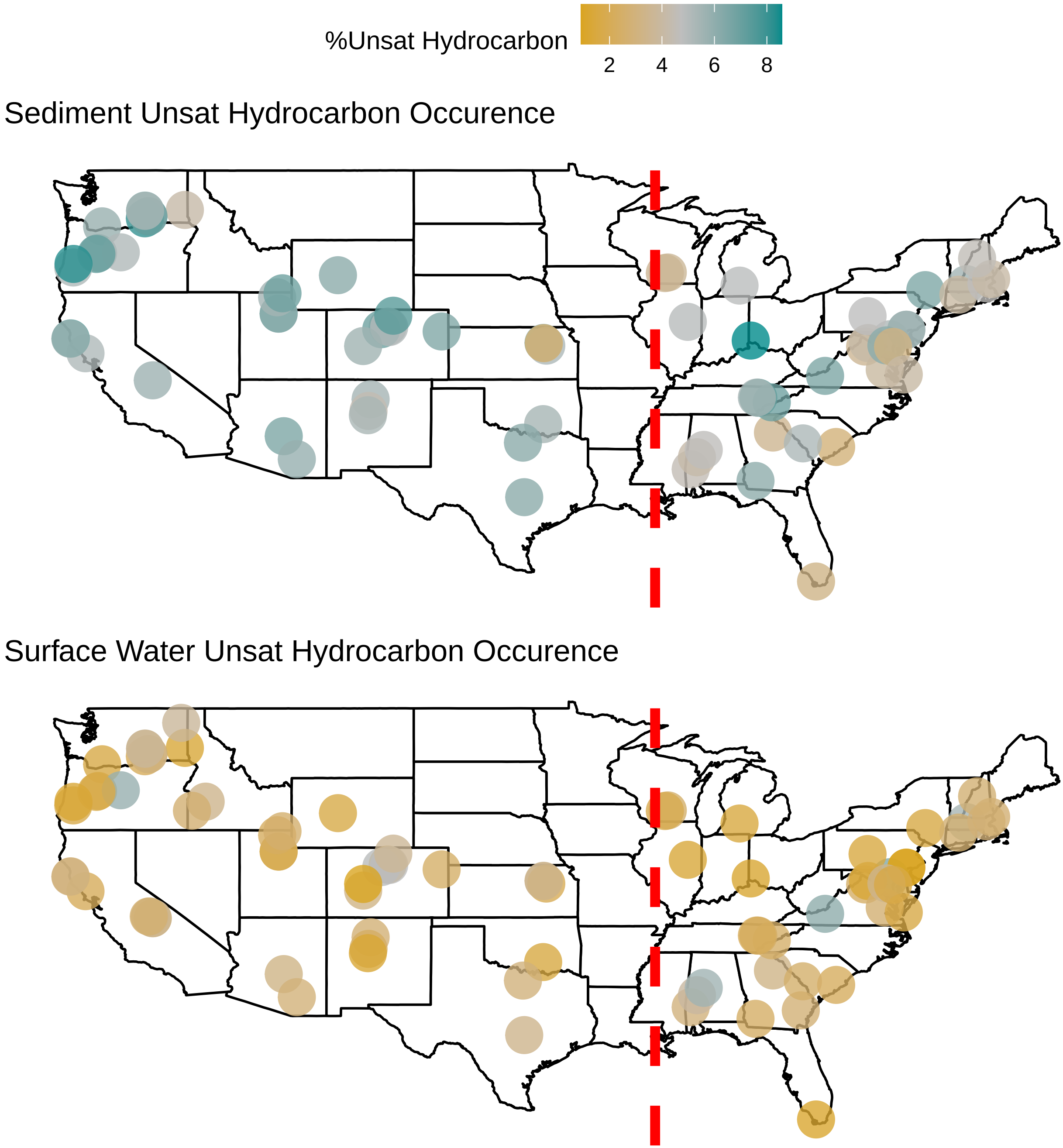

### Supplementary File 1

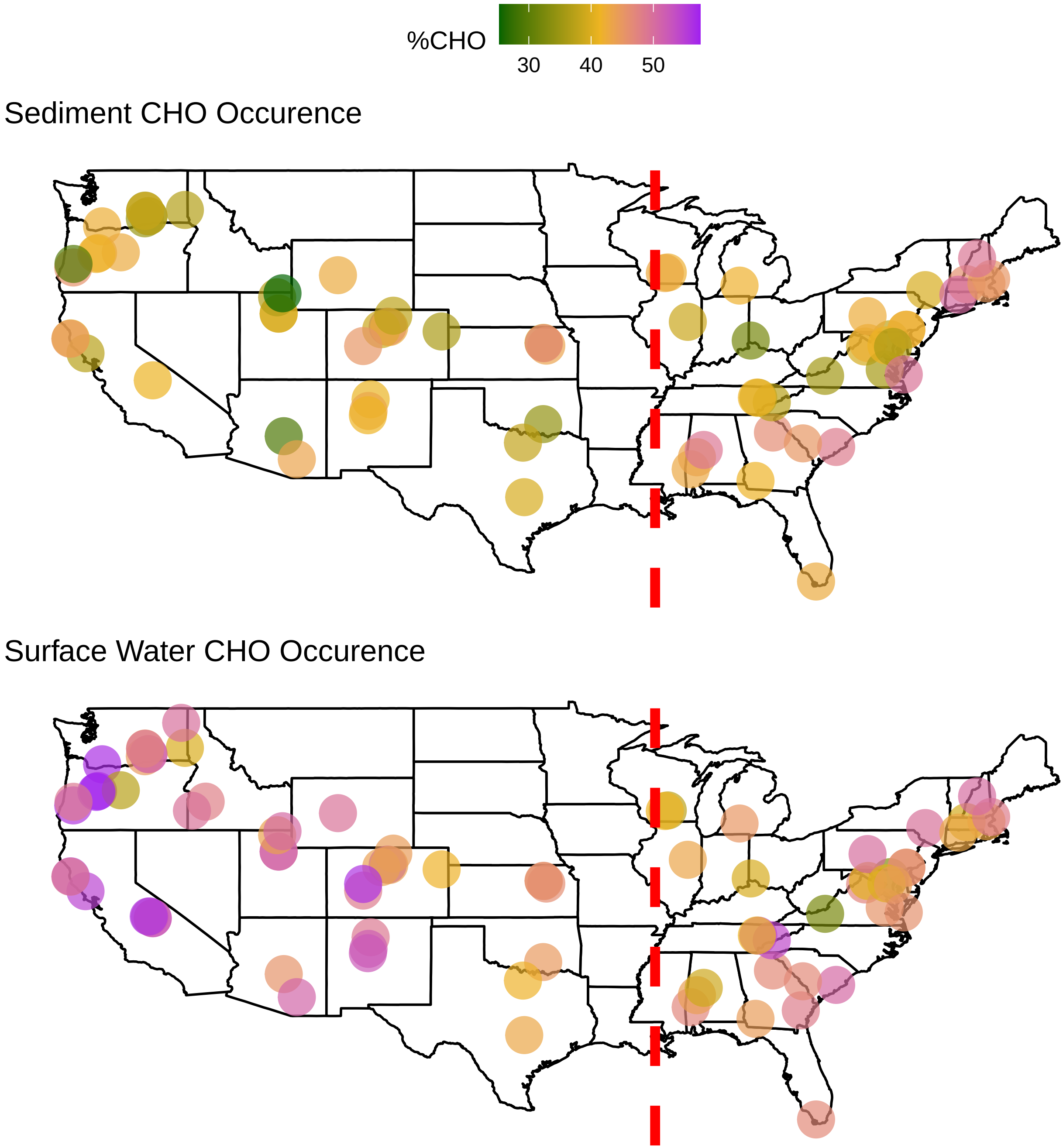

### Supplementary File 1

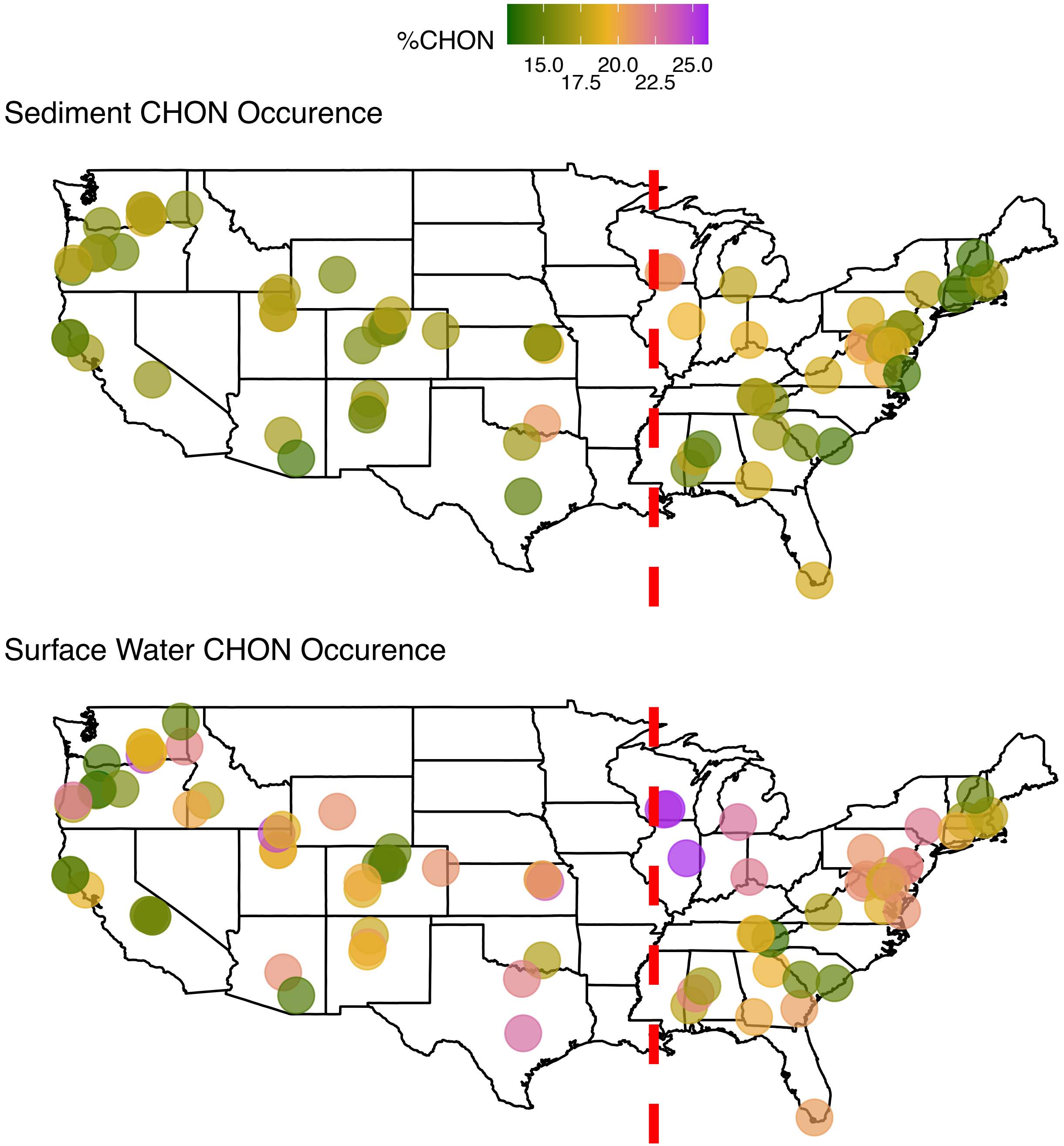

### Supplementary File 1

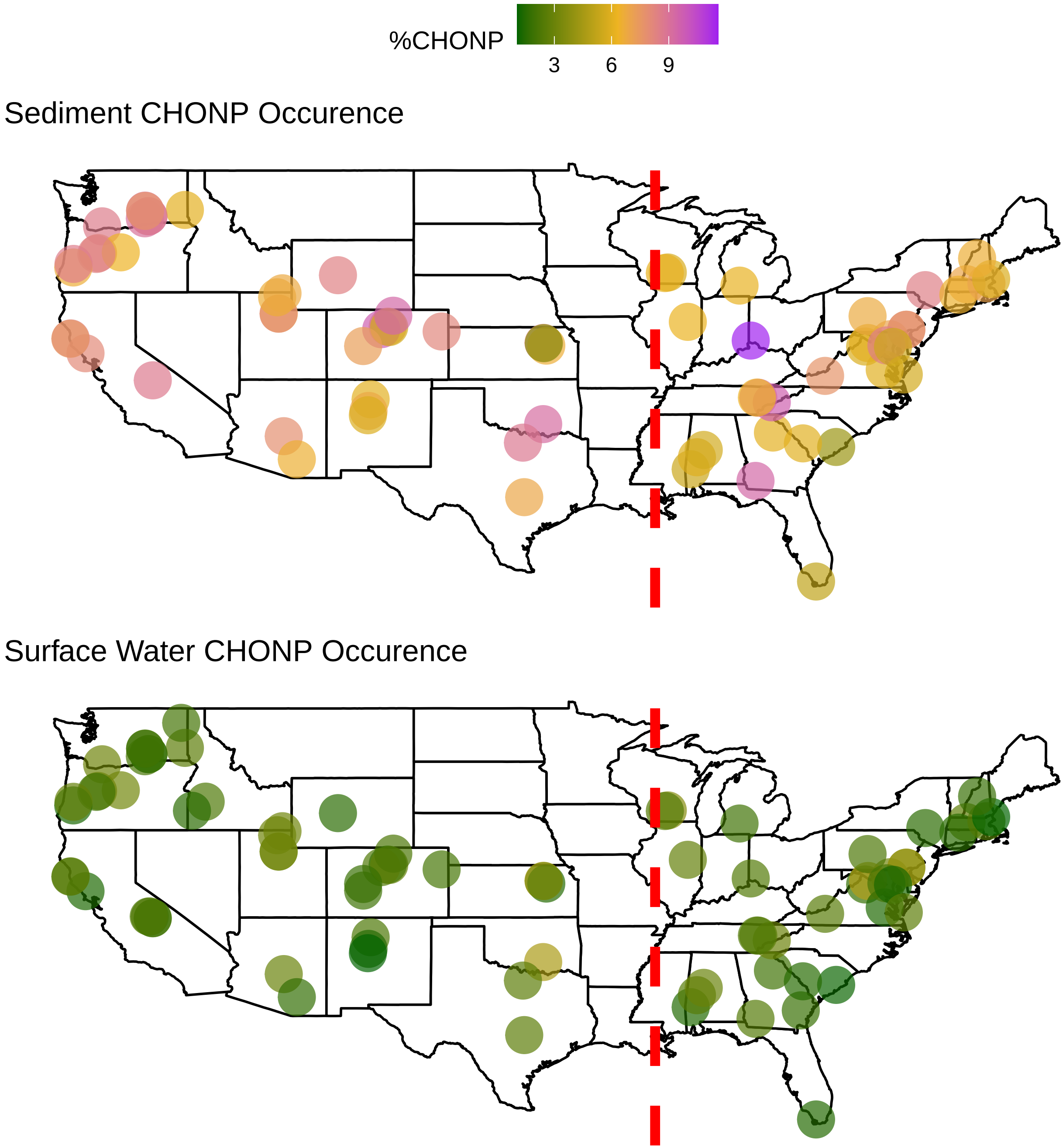

### Supplementary File 1

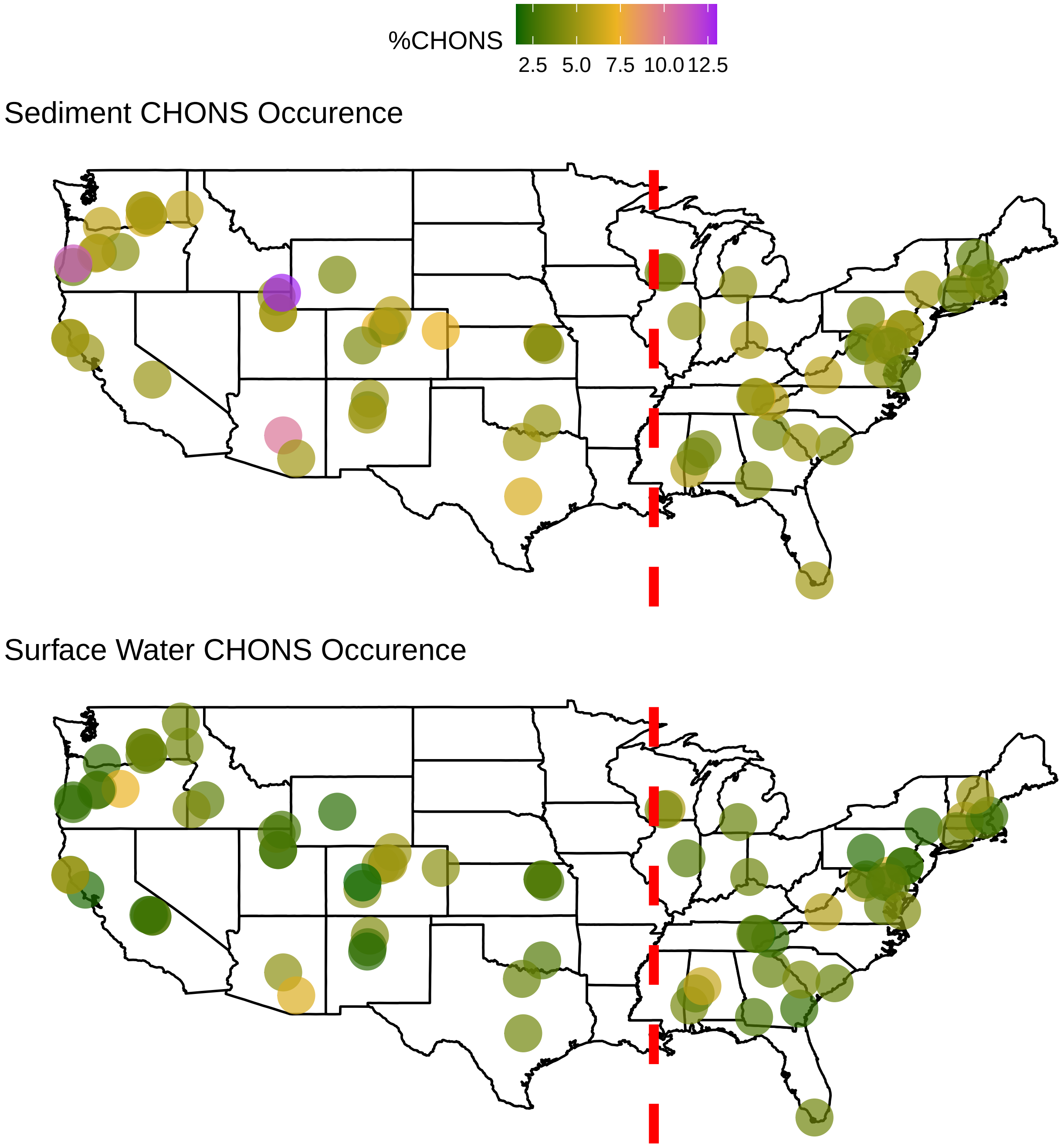

### Supplementary File 1

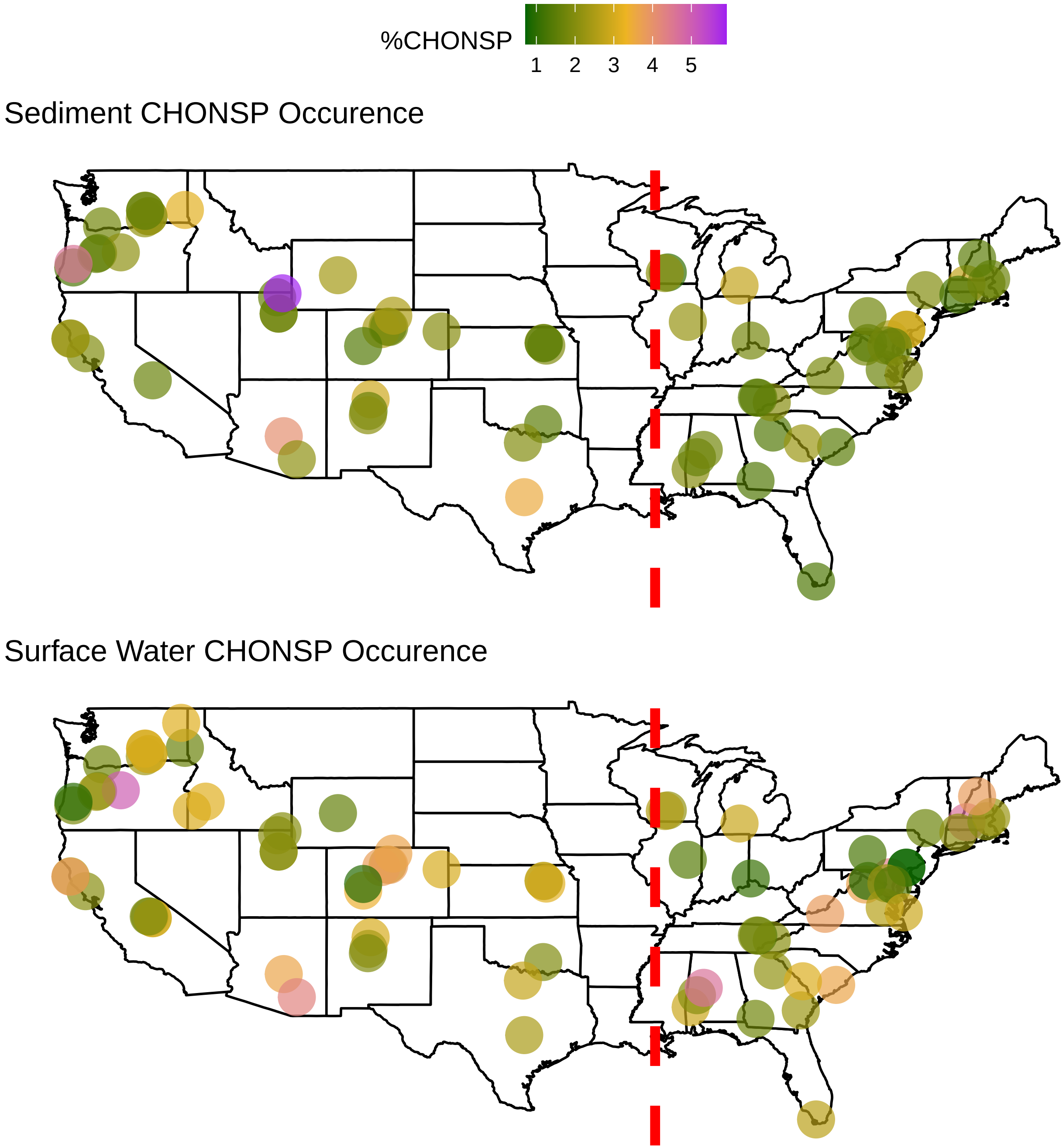

### Supplementary File 1

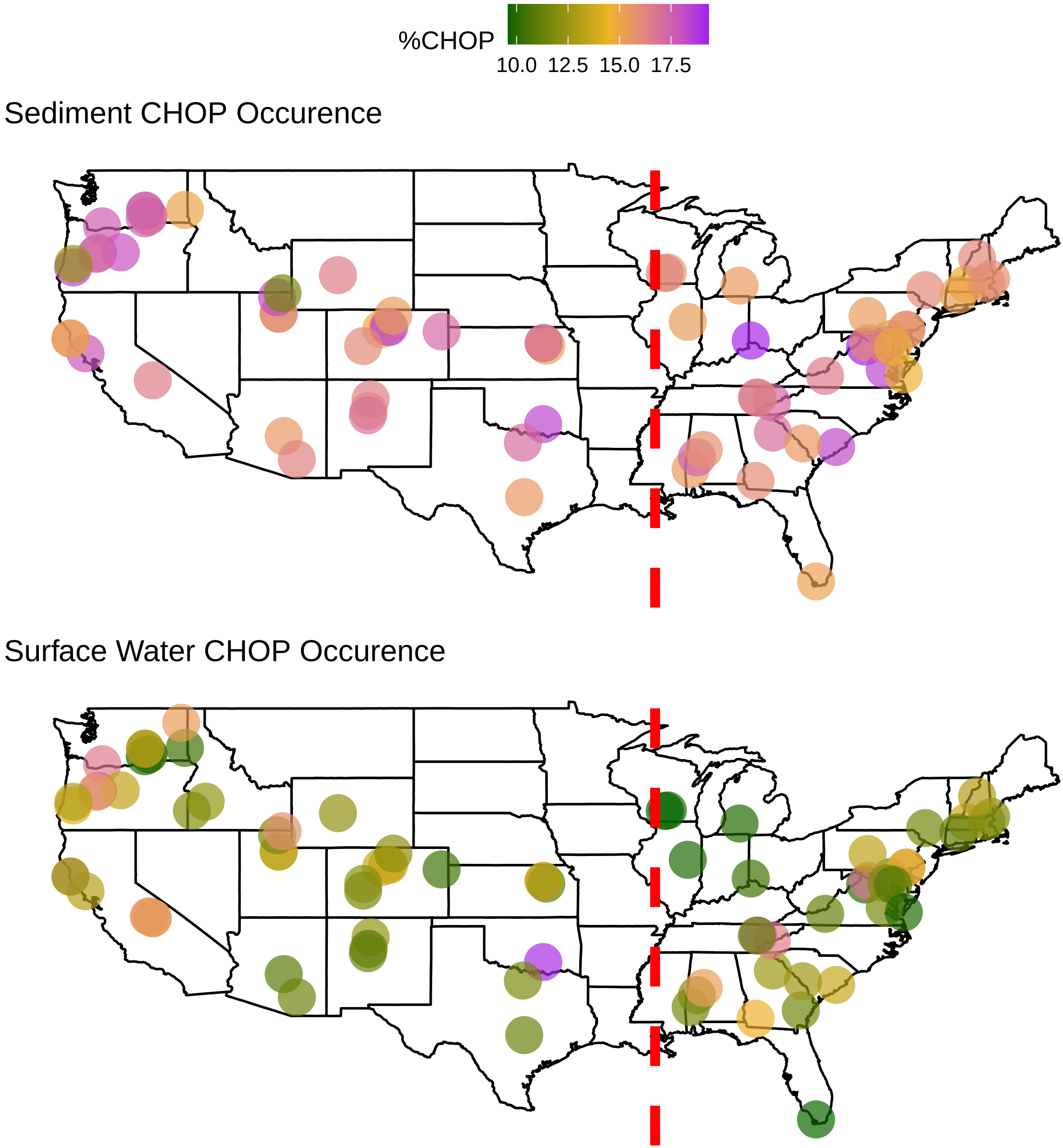

### Supplementary File 1

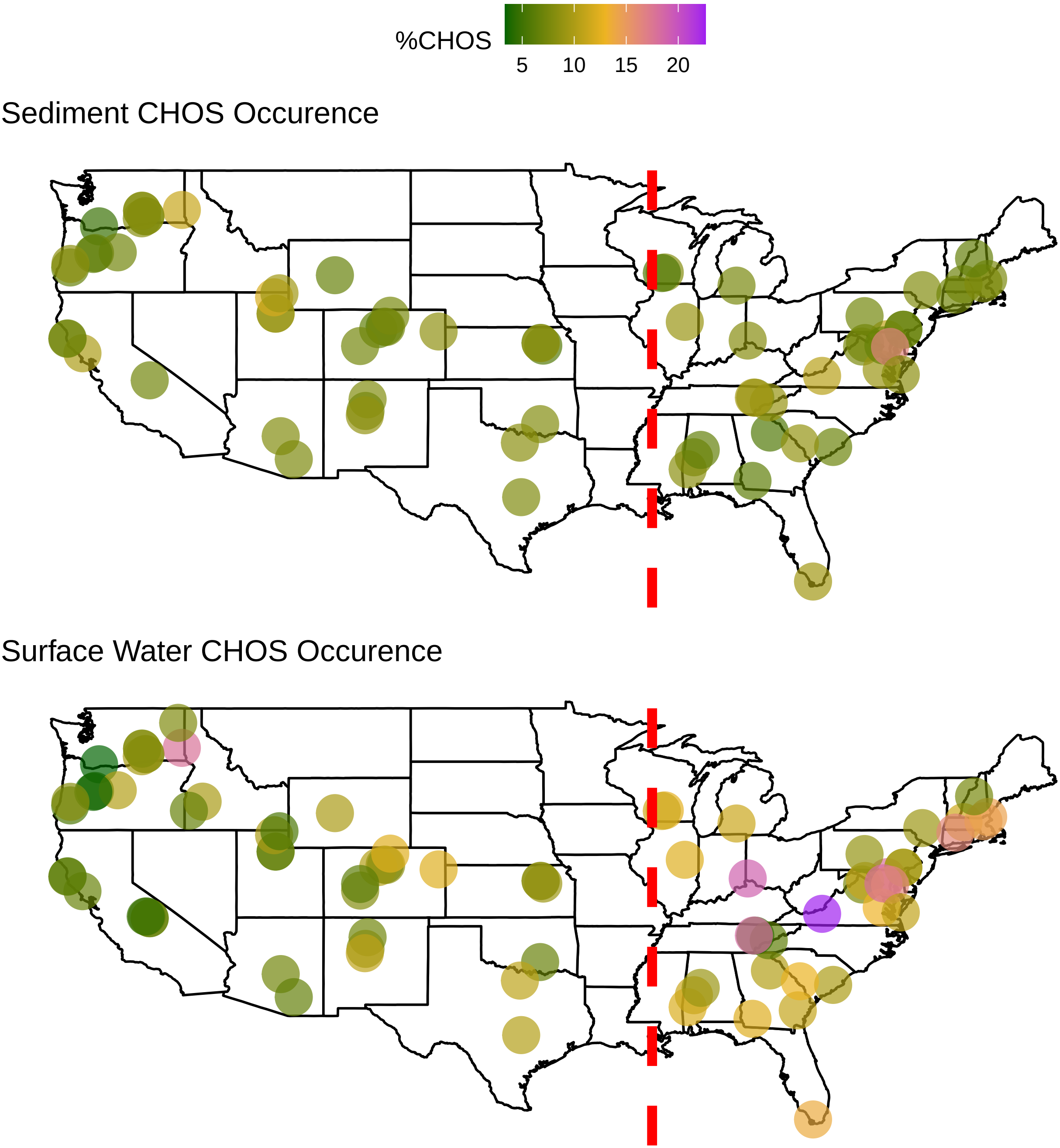

### Supplementary File 1

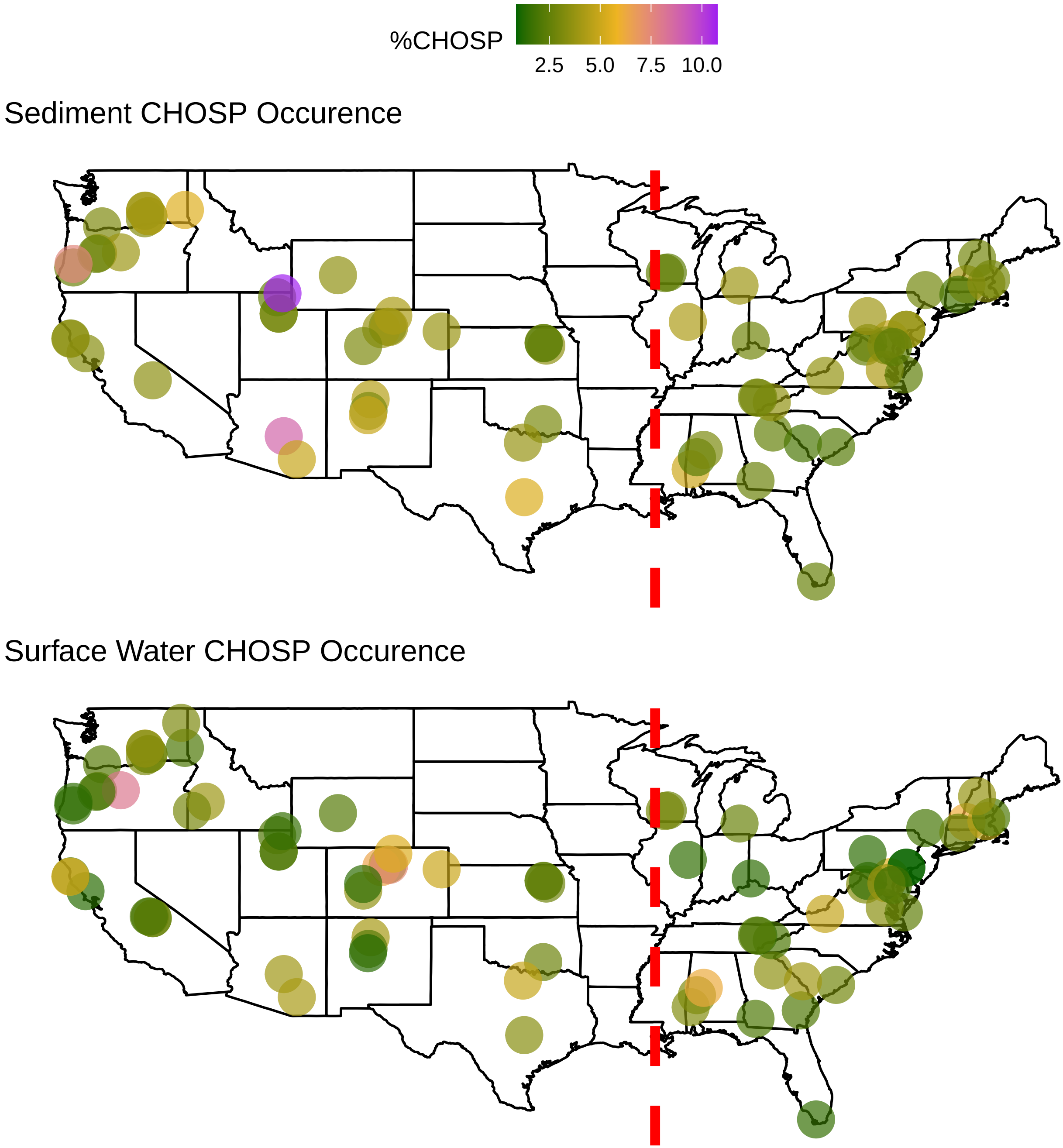

### Supplementary File 1

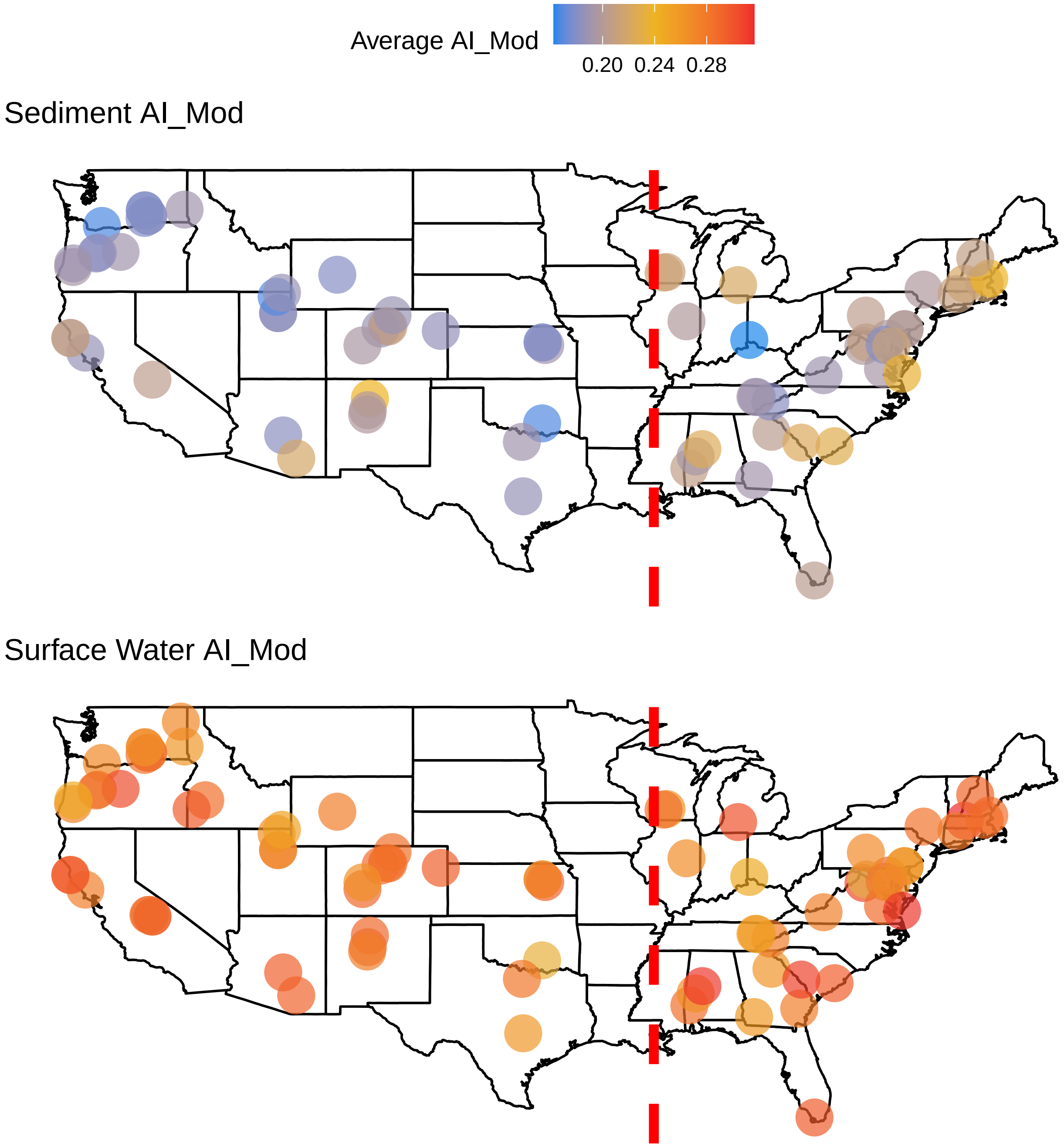

### Supplementary File 1

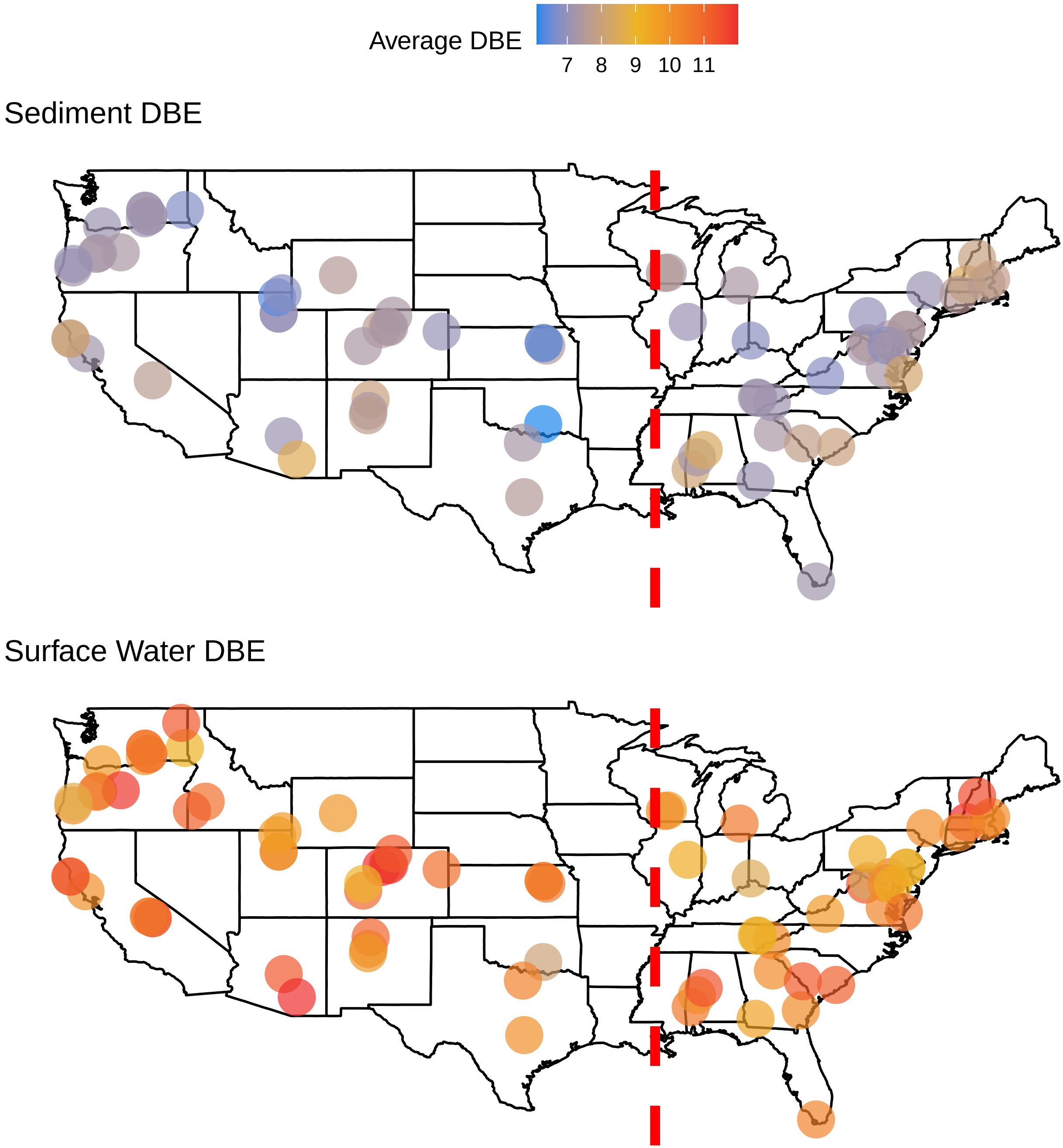

### Supplementary File 1

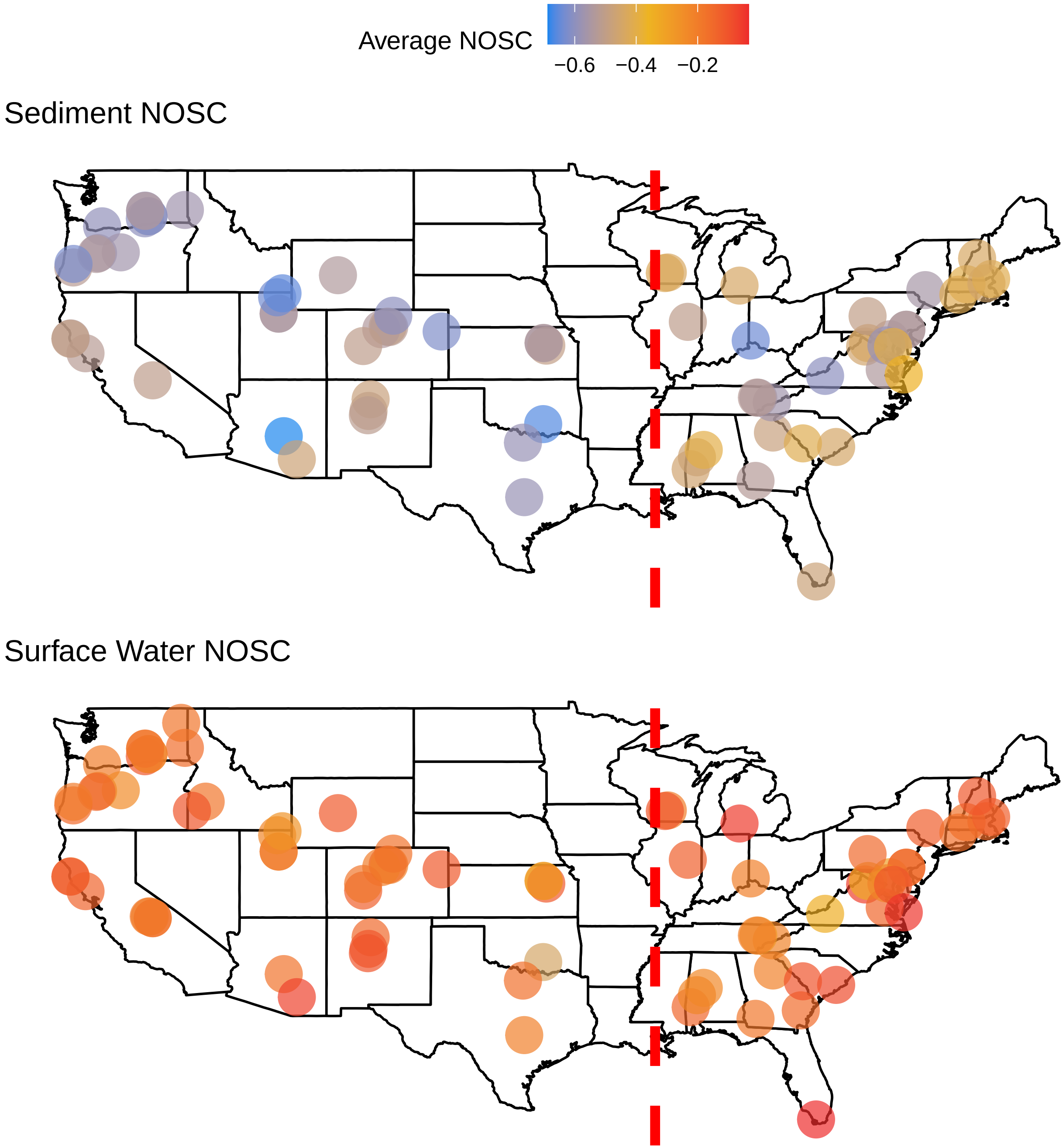

### Supplementary File 1

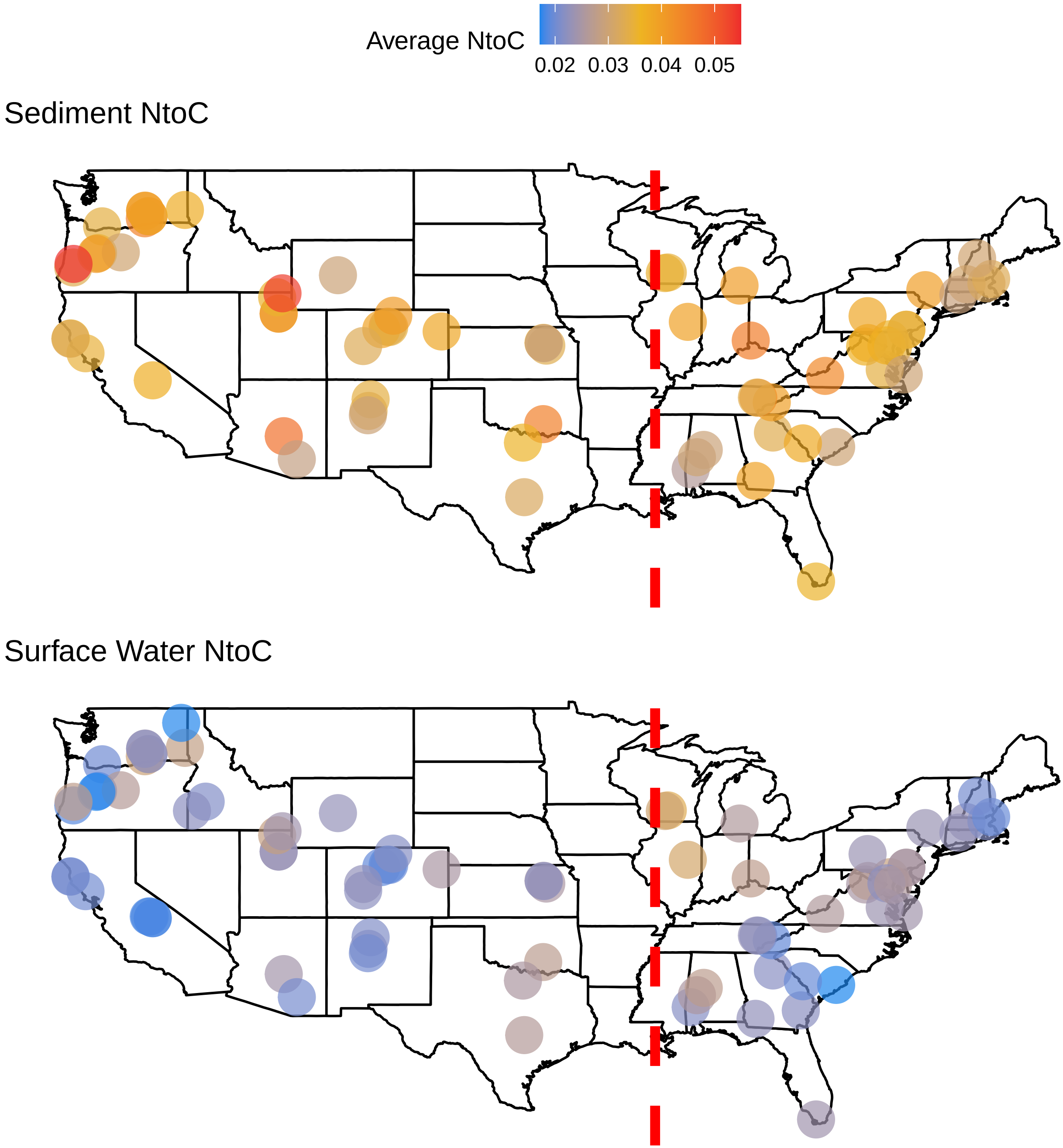

### Supplementary File 1

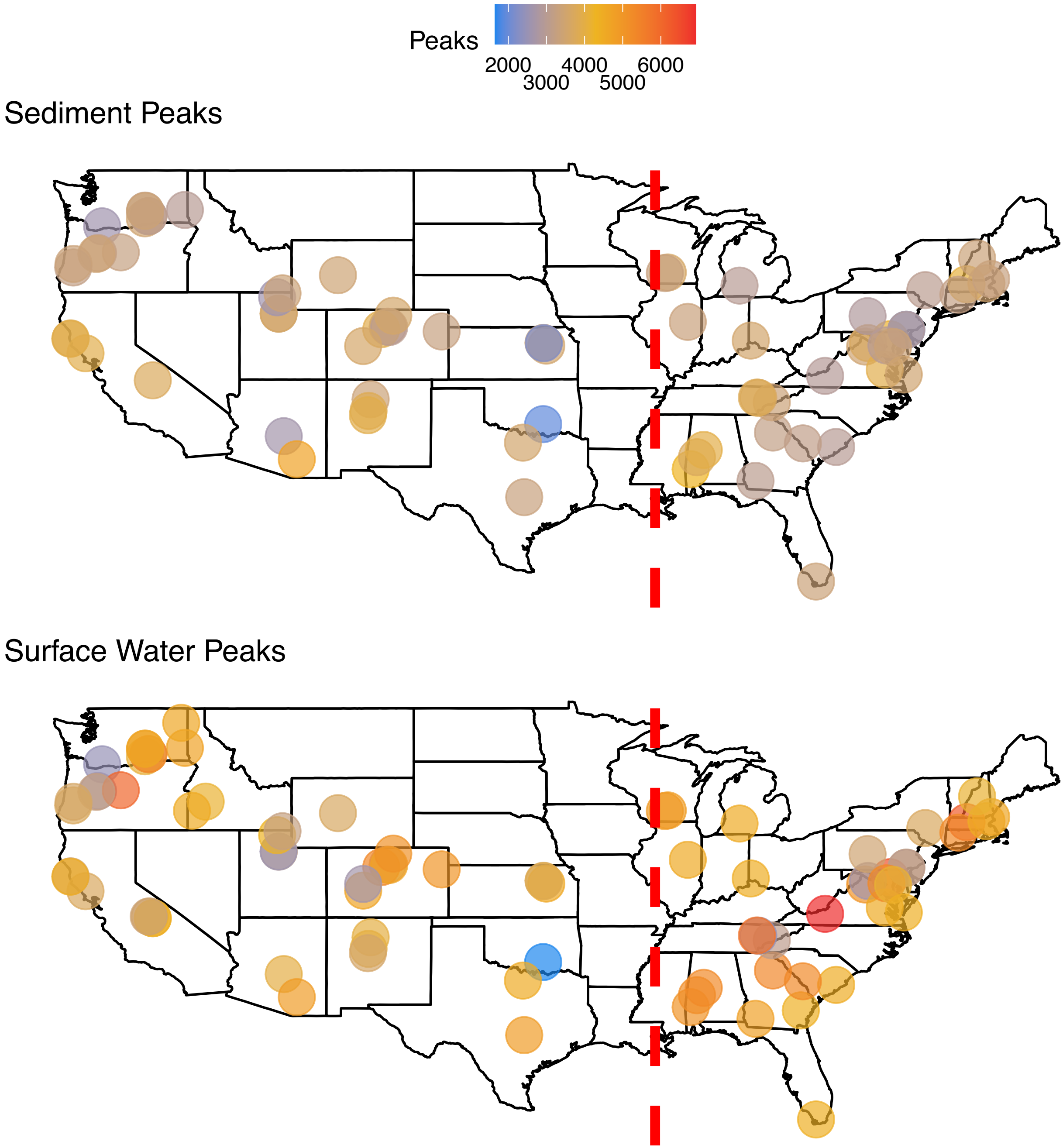

### Supplementary File 1

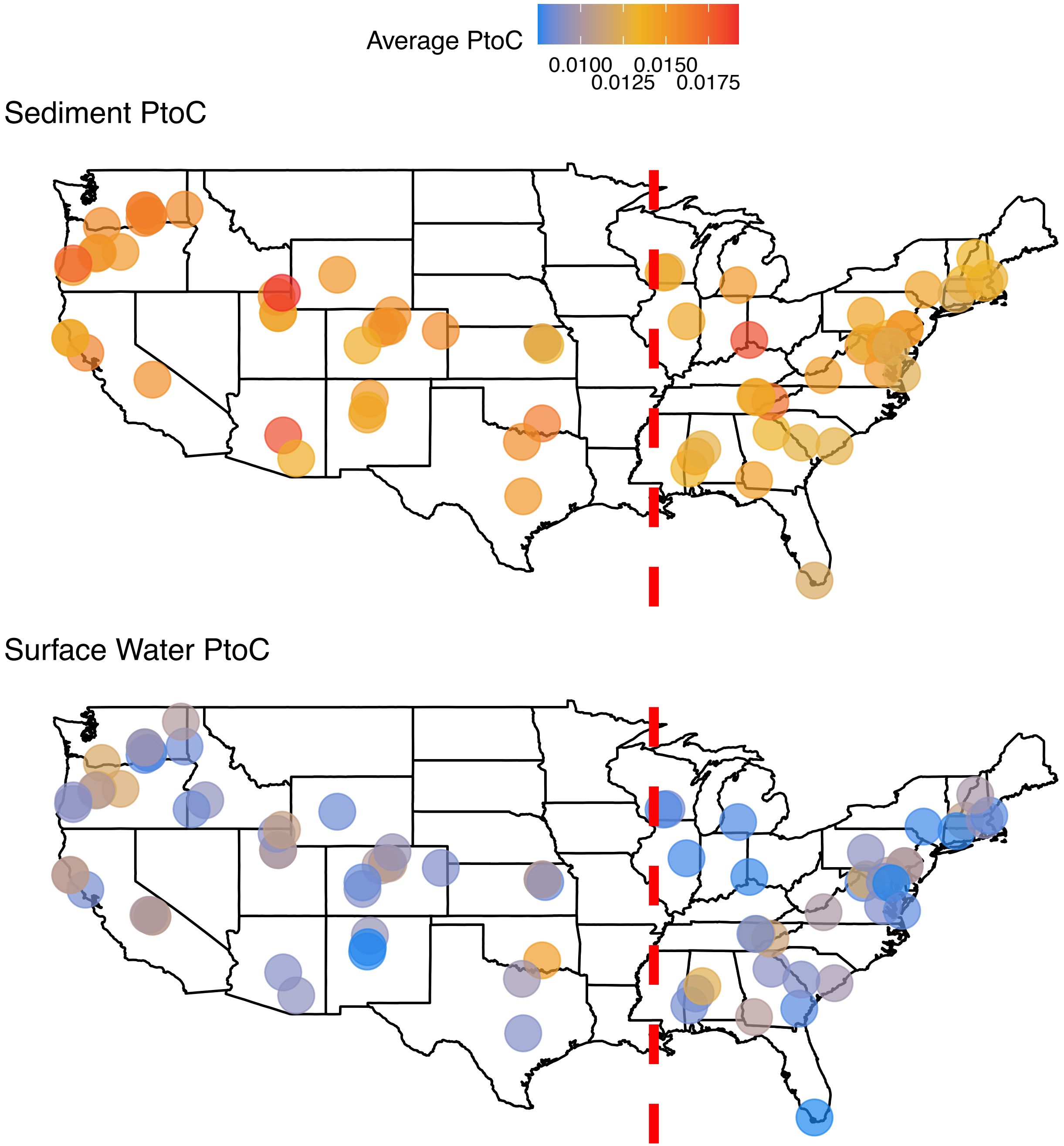

### Supplementary File 1

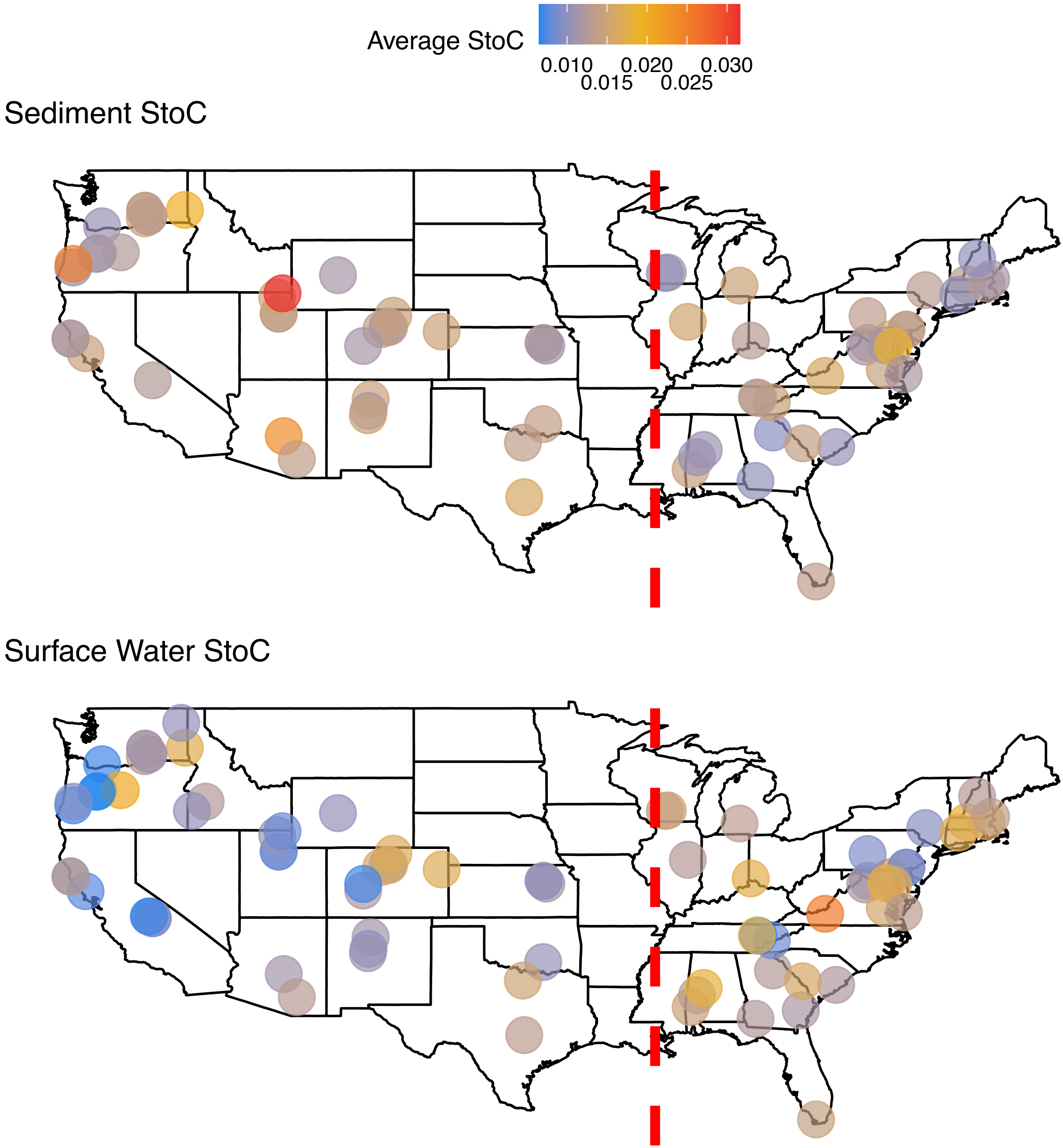

### Supplementary File 1

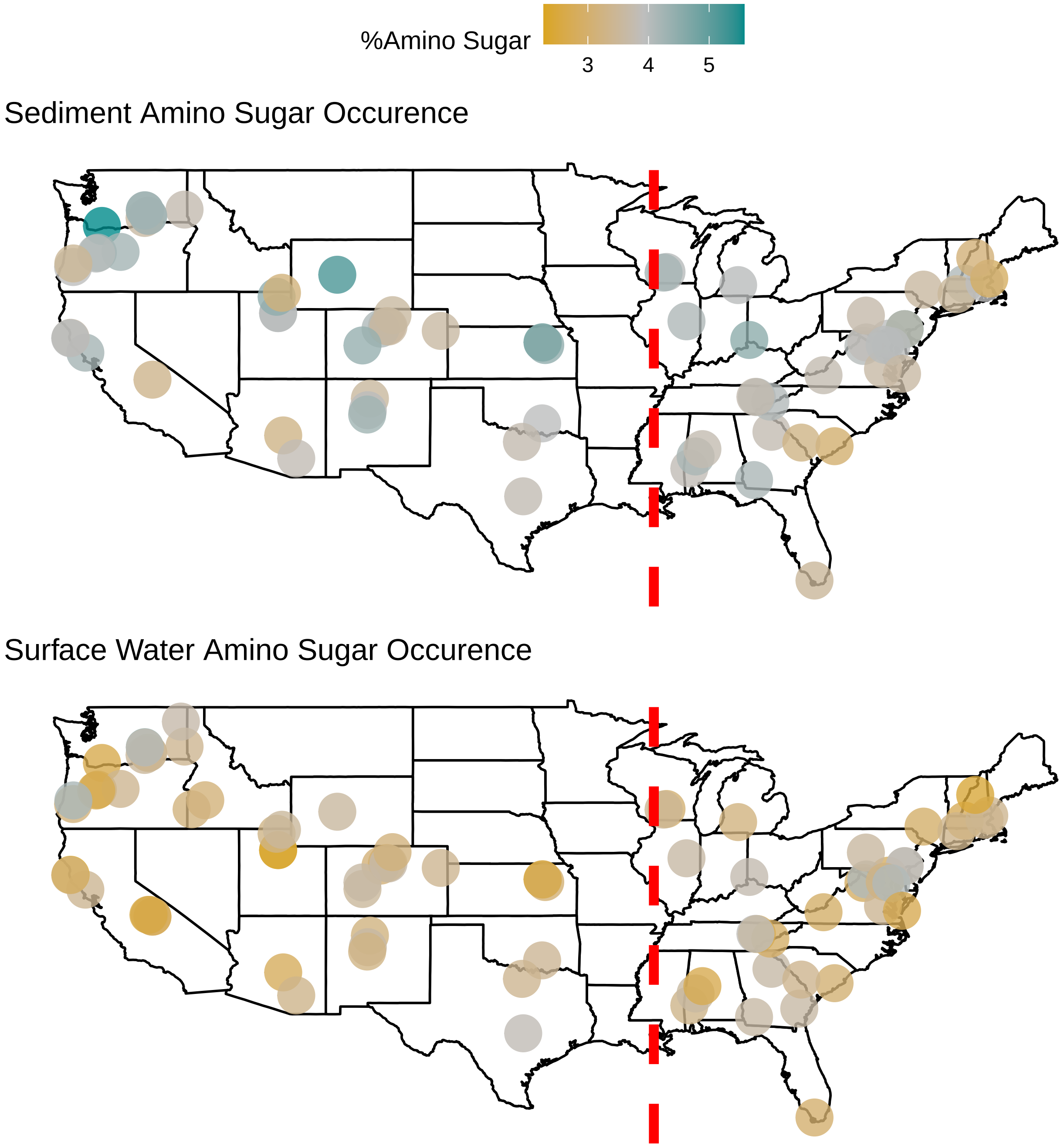

### Supplementary File 1

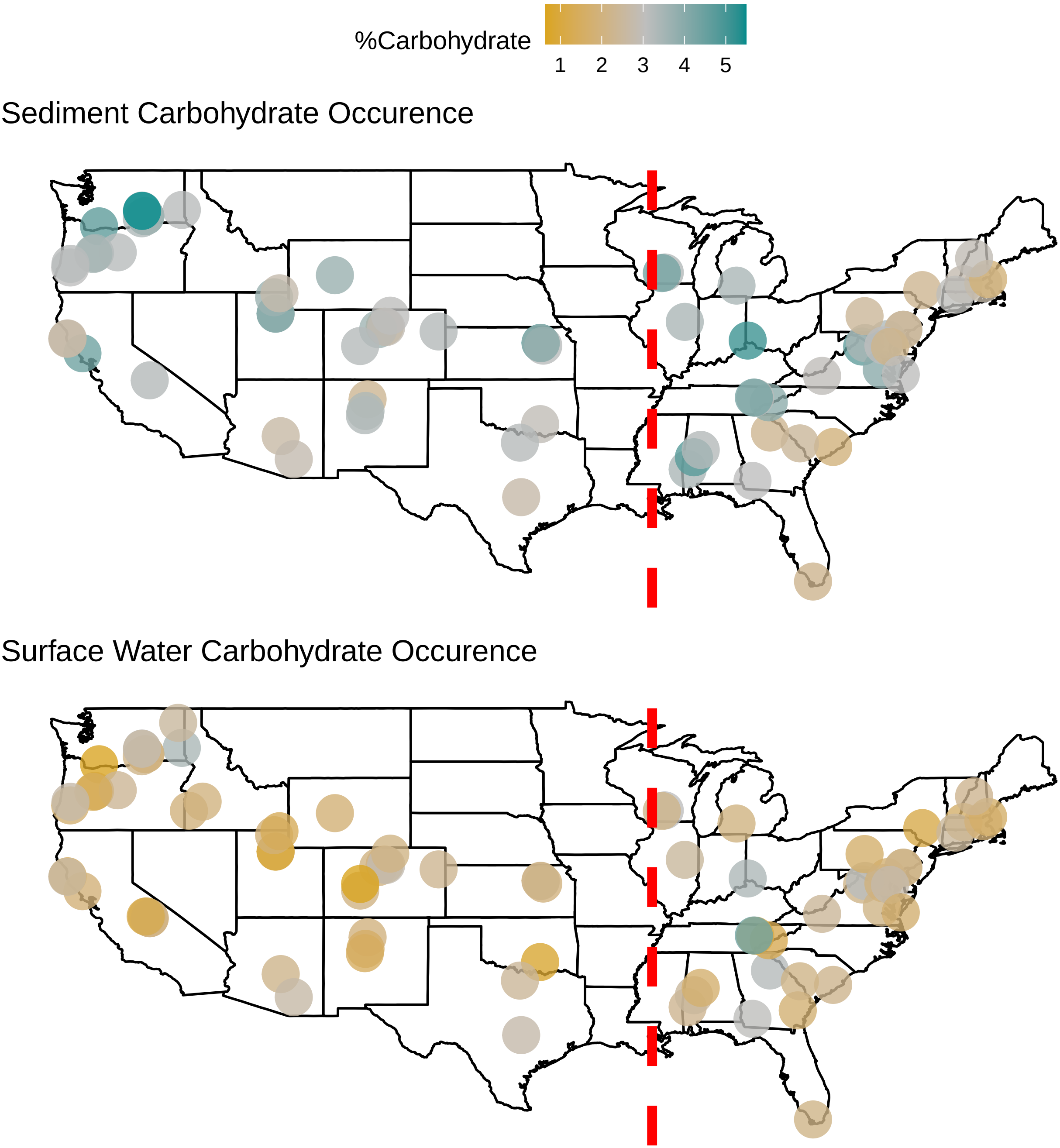

### Supplementary File 1

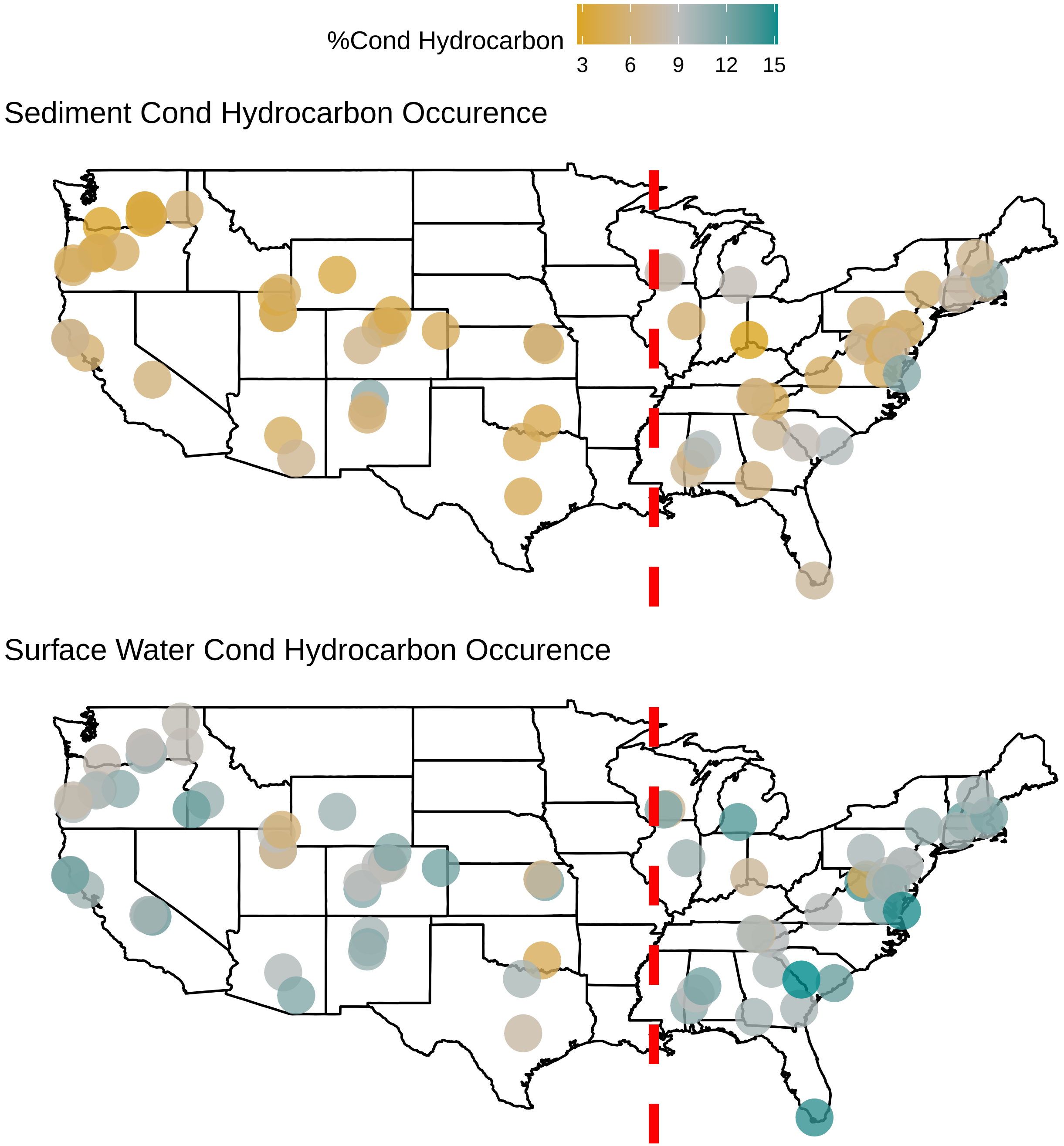
