## Supplementary File 1 for "Using Community Science to Reveal the Global Chemogeography of River Metabolomes"

**Supplemenatry_File_1_US_Maps_Properties_El_groups_Classes.zip**

To determine CONUS-scale patterns, sites were divided into eastern and western US based on their position relative to the location of the Mississippi River at St. Louis, Missouri (indicated by a red-dashed line on all maps). Replicates at each site were merged such that if a metabolite was observed in one replicate, it was considered present at the site. Maps were then generated based upon the latitude and longitude of each site (Table S2). Maps vary based upon the data that they present:

- Map names containing the word “**Metric**” feature a spatial representation of the specified derived molecular property, such that each dot is colored based on the average value of at a given site. For example, “US_Maps_Metric_NtoC.pdf” shows the average N:C ratio at each site sampled across the US.
- Map names containing the word “**El**” feature a spatial representation of a given elemental group, such that each dot represents relative abundance. For example, “US_Maps_El_CHONS.pdf” shows the percent of identified metabolites that had the elemental composition of CHONS.
- Map names containing the word “**Class**” present a spatial analysis of some compound class, such that each dot is colored based upon the relative abundance of a single class at a given site. For example, “US_Maps_Class_Tannin.pdf” shows the percent of identified metabolites at some site that were assigned to ‘tannin’ compound class.
